# pH-dependent direct sulfhydrylation pathway is required for the pathogenesis of *Mycobacterium tuberculosis*

**DOI:** 10.1101/2024.07.28.605477

**Authors:** Vaibhav Kumar Nain, Vishawjeet Barik, Manitosh Pandey, Mohit Pareek, Taruna Sharma, Rahul Pal, Shaifali Tyagi, Manish Bajpai, Prabhanjan Dwivedi, Bhishma Narayan Panda, Yashwant Kumar, Shailendra Asthana, Amit Kumar Pandey

## Abstract

Methionine is essential for the survival of *Mycobacterium tuberculosis* (*M. tuberculosis*) inside the host. However, the transsulfuration pathway, a major contributor of methionine, is dispensable for the growth of *M. tuberculosis* suggesting redundancy in the methionine biosynthesis pathway. Orthologues of MetZ_TB_ in other bacterial species are known to operate a redundant single-step methionine biosynthesis pathway called *direct sulfhydrylation.* In this study, we demonstrate that genetic disruption of the *metZ*-mediated direct sulfhydrylation pathway in *M. tuberculosis* hinders growth at low pH, an effect mitigated by methionine supplementation. Computational analyses, including *in-silico* molecular docking and molecular dynamics (MD) simulations, reveal enhanced binding of the MetZ substrate, O-succinyl homoserine (OSH), to the active site of MetZ at acidic pH. Intriguingly, despite increased intracellular ATP levels, a relative decrease in the frequency of Bedaquiline (BDQ)-induced persisters is observed in *metZ*-deficient strain, suggesting a role of direct sulfhydrylation pathway in modulating BDQ sensitivity. Finally, we demonstrated that the absence of *metZ* impedes the ability of *M. tuberculosis* to grow inside the host.

## Introduction

Tuberculosis, an ancient disease, continues to pose a serious global health challenge, negatively impacting economies and the public health systems worldwide. In 2023, *Mycobacterium tuberculosis* (*M. tuberculosis*) caused 1.8 million deaths, making it the leading cause of death from a single infectious agent [1]. A steady increase in the frequency of drug-resistant tuberculosis cases has significantly complicated the therapeutic landscape. A majority of the approved drugs target pathways that are essential for the survival of *M. tuberculosis*. Targeting biosynthesis of essential amino acids as a therapeutic strategy has demonstrated immense potential and has yielded several molecules with the potential to become a future drug [2].

Methionine plays a pivotal role in cellular function, serving as the initiator amino acid, *N-formyl-*L-methionine (*f*Met), in protein synthesis in prokaryotes as well as protein degradation in eukaryotes [3]. Methionine also acts as a precursor for biosynthesis of essential metabolites that includes cysteine, biotin and SAM (S-adenosyl methionine). Given its critical importance, inhibiting some of the essential proteins involved in the methionine biosynthesis pathway has been extensively investigated as a potential strategy to identify new drugs against tuberculosis [4-8]. The transsulfuration pathway, comprising a series of enzymatic reactions and intermediates involved in methionine biosynthesis, is largely conserved across various microbial species and has been extensively reviewed [9]. Although biosynthesis of methionine is known to be essential for the growth of *M. tuberculosis* [4, 6, 10], some genes in this pathway are dispensable, indicating potential redundancy or conditional essentiality [11-13]. Several reports suggest the existence of an alternate biochemical reaction in which the metabolite O-succinyl homoserine (OSH) is directly converted into homocysteine [12, 14]. Interestingly, instead of cysteine, host-generated hydrogen sulfide (H_2_S) acts as the S-donor in this reaction catalyzed by the enzyme O-succinyl homoserine sulfhydrylase (OSHS). The direct sulfhydrylation step allows several microbial species to bypass the two-step conversion of OSH to homocysteine via cystathionine, a crucial intermediate in the canonical methionine biosynthesis pathway [9, 15-17]. The presence of a putative OSHS encoded by *metZ* (*Rv0391*) in *M. tuberculosis* suggests the existence of a yet uncharacterized active direct sulfhydrylation pathway in *M. tuberculosis* [18].

In this study, employing genetic and computational approaches, we investigate the necessity of redundant methionine biosynthesis pathways in *M. tuberculosis*. We demonstrate that *metZ* gene is required for methionine biosynthesis under acidic conditions. Using computational approach, we further demonstrate that the binding of the substrate to the active site of the enzyme MetZ is more stable at acidic pH. Using CRISPR-mediated gene silencing, we provide evidence that *metZ*-dependent direct sulfhydrylation pathway in *M tuberculosis* functions independently of the transsulfuration pathway. The failure of *M. tuberculosis* strain, lacking both the transsulfuration and direct sulfhydrylation pathways, to grow in methionine-lacking medium further validates the redundancy of these pathways and suggests that the *metZ*-mediated direct sulfhydrylation pathway becomes conditionally essential for the pathogen to survive under acidic and *in-vivo* growth conditions. In conclusion, we report that *metZ*-mediated direct sulfhydrylation pathway is the primary driver of the methionine biosynthesis under acidic conditions and is therefore essential for the survival of *M. tuberculosis* within the host.

## Results

### MetZ is dispensable for the *in-vitro* growth of *M. tuberculosis* and possibly involved in direct sulfhydrylation pathway for methionine biosynthesis

While *metZ* has been reported to be essential for the growth of *M. tuberculosis* on cholesterol [19], we hypothesized that the *Rv0391 (metZ)* gene encodes a putative O-succinylhomoserine sulfhydrylase (OSHS) that is responsible for direct sulfhydrylation (Fig. 1A). We performed a bioinformatics-based analysis using the protein sequence of MetZ_TB_ and found that the putative sequence belonged to the cluster of orthologues group ENOG502GJ5S which is composed of 1097 protein sequences orthologous to MetZ_TB_ across 644 bacterial species. This cluster of proteins comprises various enzymes involved in cellular amino acid metabolism namely, cysteine, methionine, homocysteine, serine and aspartate. Co-incidently, all these proteins have a common PLP (pyridoxal-5-phosphate) – binding domain which is essentially a modified lysine residue that covalently binds the co-factor, hence also categorized as PLP-binding superfamily of proteins (Fig. 1B and S1A). Further, to elucidate the role of *metZ* in *M. tuberculosis*, a deletion mutant in the genetic background of H37Rv (*Rv-WT*) was generated by genetic recombineering as described previously [20]. Briefly, a hygromycin resistance cassette was inserted into the ORF encoding *metZ* (*Rv0391*) with the help of allelic exchange substrate (AES) consisting of 1kb regions of homology upstream and downstream to the target locus. The recombinants were screened for hygromycin resistance and the cassette was subsequently excised out thereby generating an unmarked mutant (Fig. 1C). The loss of the genetic sequence between 470034-471188 bases was confirmed by gene-specific PCR (Fig. S2A), DNA sequencing of the locus (Fig. S2B) and at the transcript level by RT-qPCR (Fig 1D). To check the essentiality of the *metZ* gene in growth on cholesterol, we compared the growth of *Rv-WT*, *ΔmetZ* and the complement luminescence reporter strains in different media containing glycerol (0.1%), cholesterol (200μM), propionate (10mM) as the sole carbon source. As a control, we used the *M. tuberculosis* strain lacking the *prpR* gene (*ΔprpR)* which is known to be essential for growth on cholesterol [21]. While, as expected, *ΔprpR* failed to grow on cholesterol and propionate, the growth of all the three strains were comparable irrespective to the carbon source (Figs. 1E, 1F and S3B). These findings indicate *metZ* does not contribute to the central carbon metabolism and is dispensable for *in-vitro* growth of *M. tuberculosis* on cholesterol and/ or propionate.

**Figure 1.**
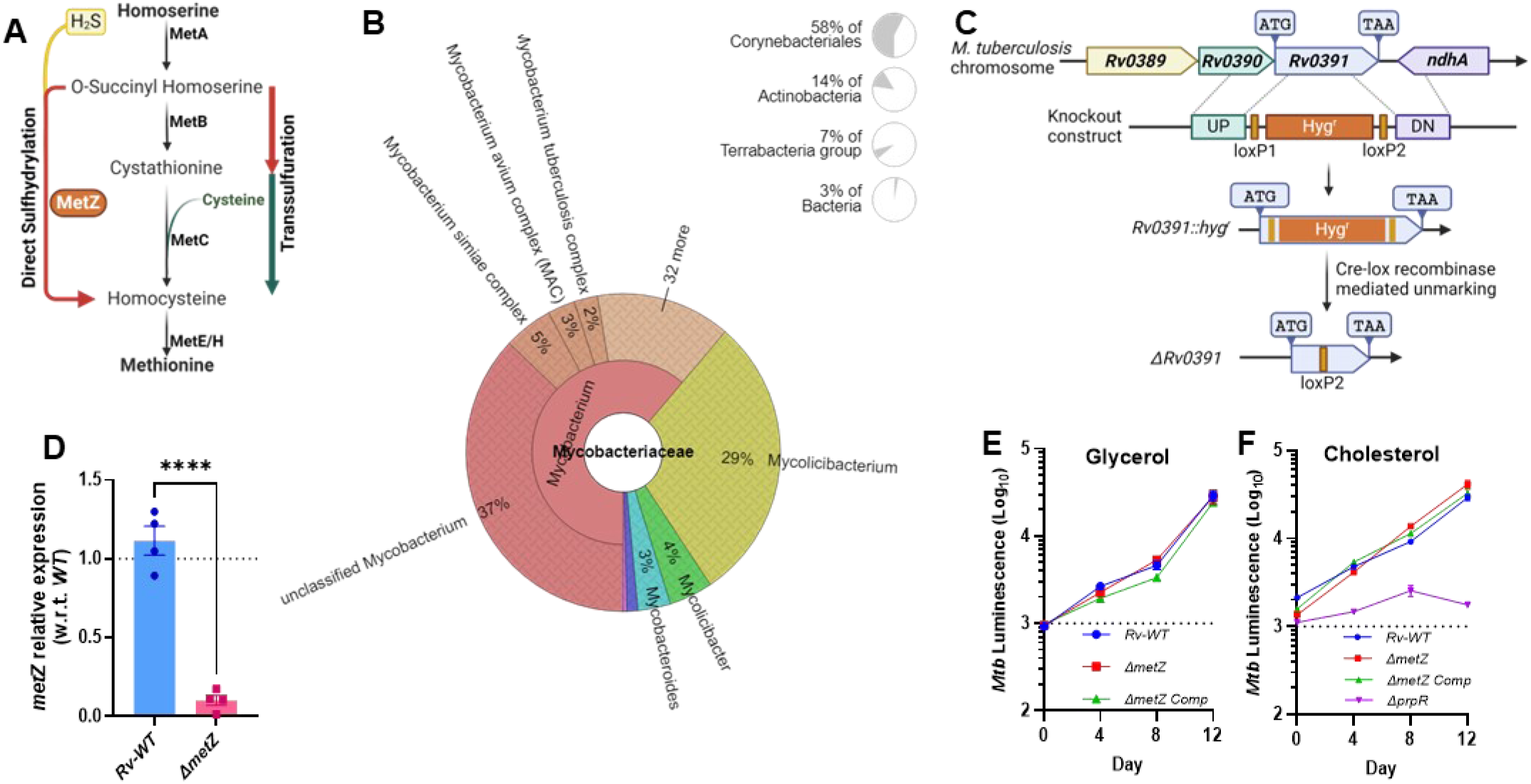
*metZ* is dispensable for growth in *M. tuberculosis.* (A). Methionine biosynthesis pathways in bacteria. Two alternate pathways exist in many bacterial species – transsulfuration where cysteine is used as an S-donor and direct sulfhydrylation wherein H_2_S is used as an S-donor. *Rv0391* encoded MetZ is such putative O-succinyl homoserine sulfhydrylase (OSHS) involved in direct sulfhydrylation in *M. tuberculosis*. (B). Cluster of Orthologues analysis for *M. tuberculosis* MetZ (Rv0391) protein sequence with member species of ENOG502GJ5S as indicated by EggNog database. (C). Schematic for knockout generation of *ΔmetZ* in *M. tuberculosis.* (D). Confirmation of *metZ* knockout using RT-qPCR for the loss of *metZ* transcript. Growth kinetics of auto-bioluminescent *ΔmetZ* with (E) glycerol (0.1%) and (F) cholesterol (200µM) as C-source. The data summary represents means ± SEM for technical replicates from two independent experiments. Statistical comparisons were made w.r.t. *Rv-WT* for (D) using Unpaired *t*-test. \*\*\*\**P*<0.0001. Comparisons not marked with statistical significance were not significant with *Rv-WT* vs *ΔmetZ* for (E) and (F).

### *metZ* deletion renders *M. tuberculosis* sensitive to physiological stressors and leads to reduced *ex-vivo* fitness

To replicate and grow inside the host, intracellular pathogens have evolved mechanisms to neutralize the host-induced physiological stressors imposed by the host environment, including oxidative stress, insult to the cell membrane, acidic pH and nutrient limitation. To study the role of *metZ* in mitigating these host-induced stress, the *M. tuberculosis* strains were exposed to a range of stress-inducing reagents. The effect of these stresses on each strain was subsequently assessed by calculating the survival index relative to the *Rv-WT* strain. In comparison to the *Rv-WT* and complement strains, *ΔmetZ,* demonstrated a reduced ability (∼40-60%) to survive under all the stress conditions tested (Fig. 2A). Further, we determined intra-bacterial redox state using respective *M. tuberculosis* strains expressing genetically encoded redox biosensor (Mrx1-roGFP2). To establish the dynamic range of the biosensor readout, *Rv-WT* and the *ΔmetZ* strains expressing Mrx-roGFP2 were treated with H_2_O_2_ and Dithiothreitol (DTT) to achieve maximum oxidation and reduction states, respectively. This defined the upper and lower limits of the biosensor response (dotted lines in Fig. 2B). We demonstrate that in comparison to *Rv-WT*, the *ΔmetZ* strain exhibited significantly higher intracellular ROS levels (Fig. 2B) suggesting an inability of the mutant to maintain redox homeostasis possibly due to impaired mycothiol recycling. Notably, unlike *Rv-WT*, we did not observe any additional increase in fluorescence signal in H_2_O_2_ treated *ΔmetZ* strain, further supporting the presence of elevated ROS levels that may have saturated the biosensor (Fig. 2B). Subsequently, increased levels of lipid hydroperoxides were detected in the *ΔmetZ,* strain compared to the *Rv-WT* and complement strains, further corroborating our previous finding that the absence of *metZ* gene in *M. tuberculosis* results in excessive accumulation of intracellular reactive oxygen species (Fig. 2C). Further, to assess the role of *metZ* in the intracellular survival of the *M. tuberculosis* inside the macrophages, resting or IFN-γ induced THP1-derived macrophages were infected with *Rv-WT*, *ΔmetZ* and the complement strains and survival indices (w.r.t. *Rv-WT*) were determined by enumerating the bacterial load by CFU (Colony Forming Unit) plating of the infected cell lysates at day 5 post-infection. We observed a significant reduction in bacterial burden in the *ΔmetZ* strain compared to both *Rv-WT* and complement strains in both resting (∼60%) and activated (∼48%) macrophages (Fig. 2D and S4A). Since restricting the phagosome-lysosome fusion is a common strategy employed by the pathogen to enhance its intracellular survival, we further investigated the role of *metZ* in regulating this process. Briefly, IFN-γ induced THP1 macrophages infected with mEmerald (GFP) expressing *Rv-WT*, *ΔmetZ* and complement strains were subsequently stained with Lysotracker. The study revealed that in comparison to the *Rv-WT* and complement strains, the *ΔmetZ* strain exhibited a higher propensity to localize within the acidic compartments of the cell (Fig. 2E and S5A). Overall, these data suggest that absence of *metZ* significantly weakens the ability of *M. tuberculosis* to resist host-induced physiological stresses and evade phago-lysosome fusion, both of which are crucial for the survival of the pathogen within the host cell.

**Figure 2:**
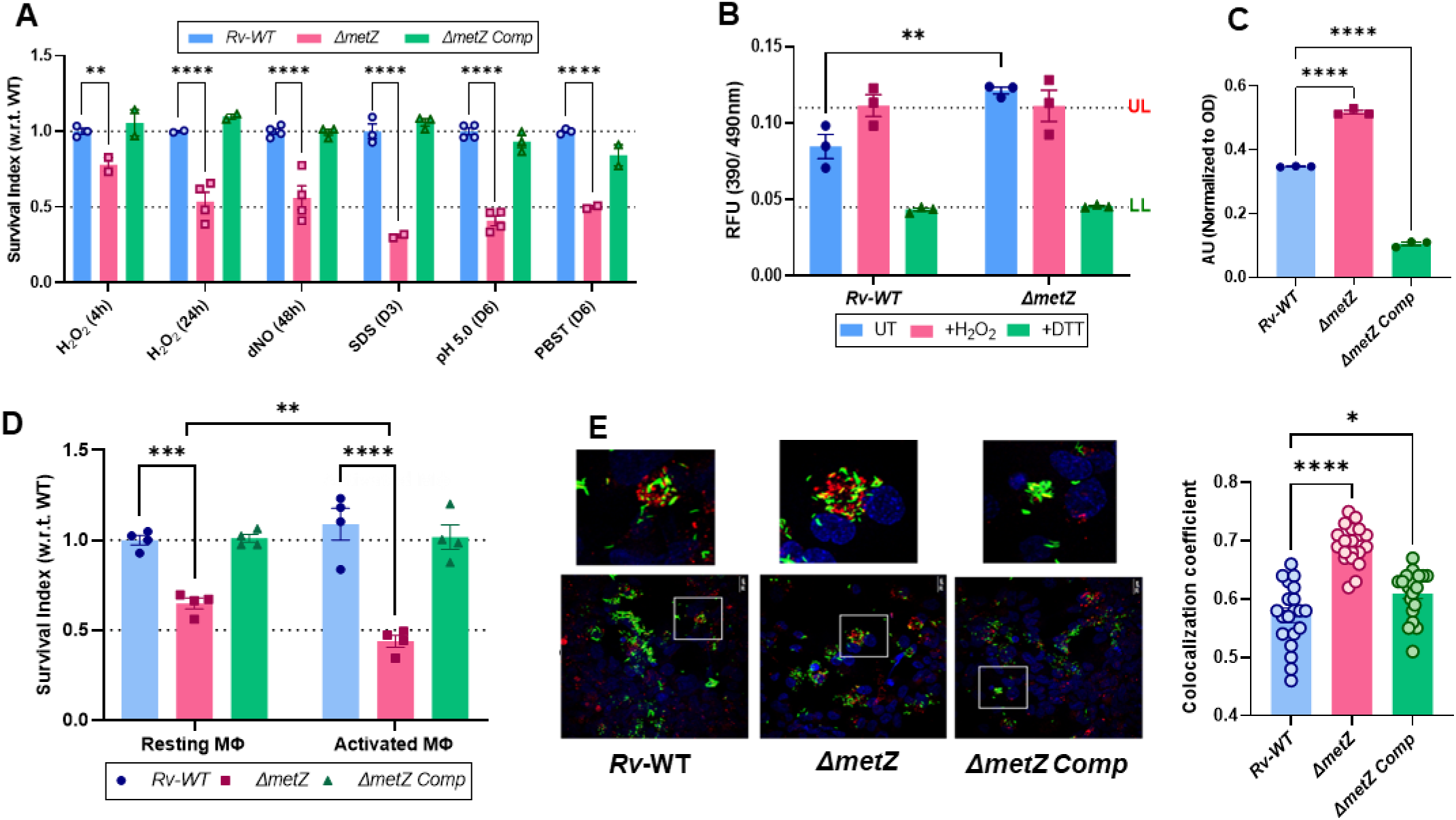
*metZ* deletion renders *M. tuberculosis* sensitive to physiological stressors and reduced *ex-vivo* fitness. (A). *In-vitro* stress survival assay. Mycobacterial cultures were treated with reagents mimicking host-induced stresses: H_2_O_2_ (Oxidative stress), DetaNO (Nitrosative stress), SDS (Surfactant stress), low pH (5.0), PSBT (nutrient starvation) *in-vitro* and survival was determined by CFU plating. (B). Intrabacterial redox state determination of *ΔmetZ* compared to *Rv-WT* using genetically encoded redox biosensor Mrx1-roGFP2. (C). Lipid hydroperoxidation levels in *ΔmetZ* compared to *Rv-WT* and complement strains using FOX2 colorimetry assay. (D). Intra-macrophage survival assay using THP-1 monocyte-derived macrophages resting or activated using IFN-γ. The survival index was determined by CFU enumeration. (E). Intracellular co-localization of GFP labelled mycobacterial strains with LysoTracker stained acidic compartments inside IFN-γ activated macrophages. Data is represented as mean ± SEM of the replicates. Comparisons for statistical significance have been made with *Rv-WT* (Control) using two-way ANOVA (Dunnett’s multiple comparisons test) for (A) where **P = 0.0084, ****P<0.0001; two-way ANOVA (Sidak’s multiple comparisons test) for (B) where **P = 0.0034; ordinary one-way ANOVA (Dunnett’s multiple comparisons test) for (C) where ****P<0.0001; two-way ANOVA (Dunnett’s multiple comparisons test) for (D) where ***P = 0.0002, ****P<0.0001; comparison of *ΔmetZ* (Resting vs. Activated macrophages) using unpaired t-test for (D) where **P = 0.0034 and ordinary one-way ANOVA for (E) where, *P = 0.0166, ****P<0.0001.

### Direct sulfhydrylation pathway is essential for *M. tuberculosis* to survive under low pH

The observed data suggest that the *ΔmetZ* strain might exhibit increased sensitivity on exposure to an acidic environment. To investigate this, we cultured different bacterial strains (*Rv-WT*, *ΔmetZ* and complement) at varying pH values and monitored their growth kinetics. While the growth kinetics were comparable at neutral pH (7.2) for all the strains, a decrease in pH resulted in a discernible growth defect in the *ΔmetZ* strain, ranging from observable impairment at pH 5.5 to complete growth arrest at pH 5.0, relative to the *Rv-WT* and complement strain (Fig. 3A and S6A and B). To further understand the impact of acidic pH on the expression of genes involved in transsulfuration (*metB* and *metC*) and direct sulfhydrylation (*metZ*), we quantified the relative expression of these genes in *M. tuberculosis* cultured at pH 7.2 and pH 5.5. RT-qPCR analysis revealed an ∼12-fold increase in the expression of *metZ* gene at pH 5.5 compared to pH 7.2, suggesting transcriptional induction of the *metZ* gene under acidic conditions (Fig. 3B). To differentiate between bacterial killing and metabolic slowdown as the cause of reduced CFU counts in the *ΔmetZ* strain at pH 5.5 (Fig. S6C), we assessed metabolic activity using Resazurin dye [22]. We found that, in contrast to the *Rv-WT* and complement strains, the *ΔmetZ* strain exhibited significantly reduced metabolic activity at pH 5.5, as indicated by lack of color change and low fluorescence, suggesting a significant impairment in metabolic function (Fig. 3C). Surprisingly, despite this metabolic impairment, ATP levels in the *ΔmetZ* strain were found to be higher than those in the *Rv-WT* and complement strains when cultured at acidic pH 5.5 (Fig. 3D).

**Figure 3:**
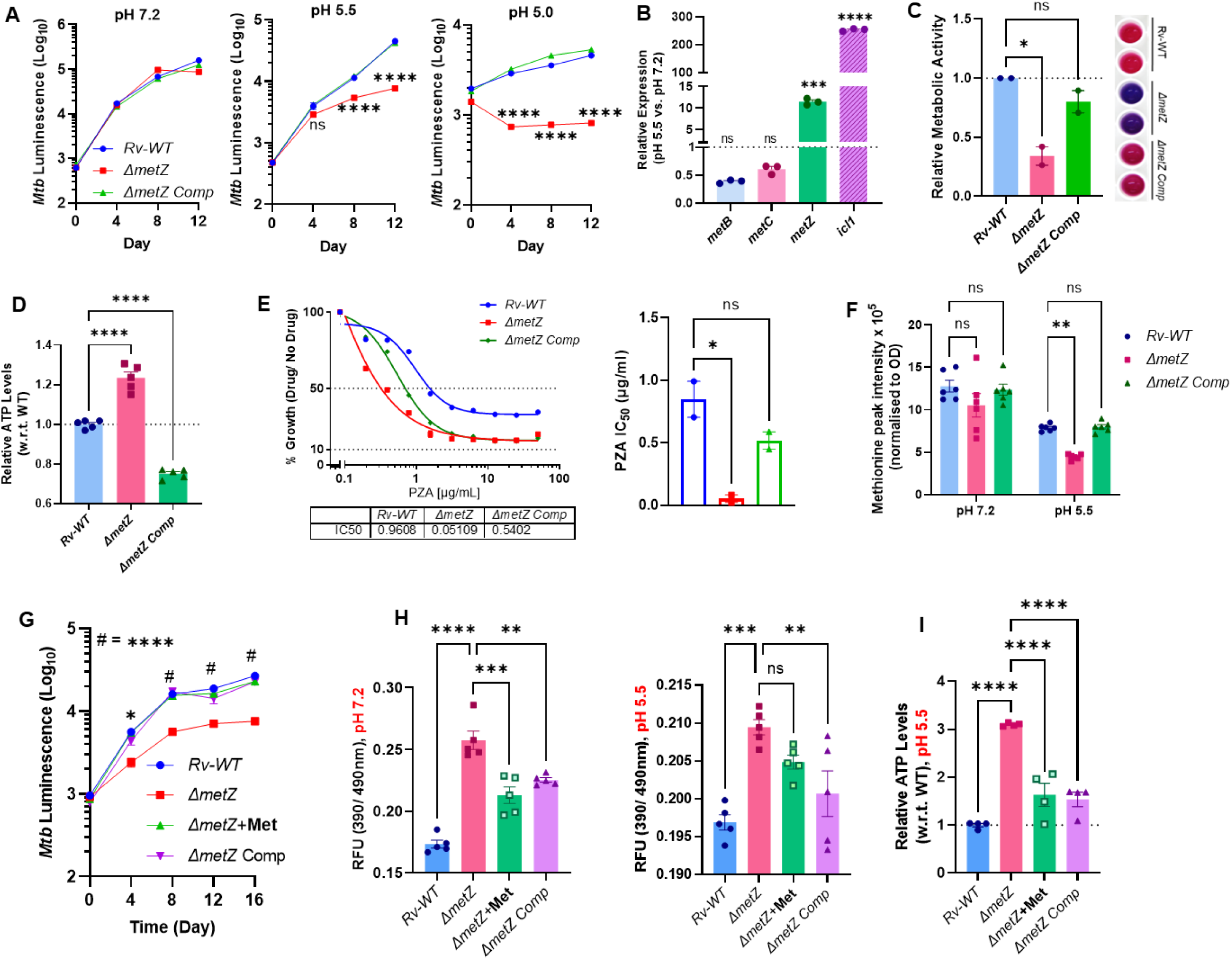
Direct sulfhydrylation pathway is essential for *M. tuberculosis* to survive at low pH. (A) *In-vitro* growth kinetics of *ΔmetZ* at various pH – 7.2, 5.5 and 5.0. (B). Expression analysis of genes involved in transsulfuration (*metB* and *metC*) and direct sulfhydrylation (*metZ*) using RT-qPCR. (C). Metabolic activity of *ΔmetZ* at acidic pH using redox dye AlamarBlue Assay. (D). Intrabacterial ATP levels estimation of *ΔmetZ* at acidic pH 5.5. (E). Pyrazinamide (PZA) sensitivity assay of *ΔmetZ* at pH 5.5. IC50 of PZA for the respective strains at acidic pH 5.5 was determined and represented as a dose-response curve. (F). Methionine estimation at pH 7.2 and pH 5.5 was performed using LC-MS. (G). Growth defect in *ΔmetZ* at pH 5.5 was rescued using exogenous methionine (Met) supplementation. (H). The intrabacterial redox state of *ΔmetZ* was determined using the Mrx1-roGFP2 redox biosensor upon supplementation with methionine (Met) at pH 7.2 and 5.5. (I). Bacterial ATP levels at pH 5.5, upon supplementation of methionine (Met). The dose-response curve for PZA was made by Non-linear regression analysis using GraphPad Prism. Data is represented as mean ± SEM of the replicates. Comparisons for statistical significance have been made using two-way ANOVA (Dunnett’s multiple comparisons test) for (A) where, ****P<0.0001, ns = non-significant; ordinary one-way ANOVA (Dunnett’s multiple comparisons test) for (B) where, ***P = 0.004, ****P<0.0001, ns = non-significant; ordinary one-way ANOVA (Dunnett’s multiple comparisons test) for (C) where *P = 0.011, ns = non-significant; ordinary one-way ANOVA (Dunnett’s multiple comparisons test) for (D) where ****P<0.0001; ordinary one-way ANOVA (Dunnett’s multiple comparisons test) for (E) where *P=0.0155, ns = non-significant; two-way ANOVA (Dunnett’s multiple comparisons test) for (F) where **P = 0.0030, ns = non-significant; two-way ANOVA (Dunnett’s multiple comparisons test) for (G) w.r.t. *ΔmetZ* where *P=0.0157, # = ****P<0.0001; ordinary one-way ANOVA (Dunnett’s multiple comparisons test) for (H) w.r.t. *ΔmetZ* where **P<0.006, ***P<0.0004, ****P<0.0001, ns = non-significant; ordinary one-way ANOVA (Dunnett’s multiple comparisons test) for (I) w.r.t. *ΔmetZ* where ****P<0.0001.

Given that the first-line anti-TB drug - Pyrazinamide (PZA) is active only at acidic pH and efficiently kills slow or non-replicating cells, we hypothesized that the differences in the cytoplasmic pH will correlate with PZA sensitivity. As expected, we observed a ∼19-fold and 4-fold increase in the sensitivity of the *ΔmetZ* strain to PZA treatment compared to *Rv-WT* and complement strain, respectively (Fig. 3E). To confirm, if the observed phenotype is a direct consequence of a defect in the methionine biosynthesis, we quantified the intracellular methionine levels. We found significantly lower levels of methionine in both *Rv-WT* and *ΔmetZ* strains grown in acidic pH 5.5 compared to those grown at pH 7.2 (Fig. 3F). Exogenous supplementation of methionine rescued the growth defect phenotype of *ΔmetZ* observed at pH 5.5 (Fig. 3G). Importantly, the addition of methionine did not alter the pH of the culture media (Fig. S6D and E). Furthermore, we observed a reduction in the intracellular ROS levels in *ΔmetZ* upon exogenous supply of methionine at pH 7.2 while this effect was less pronounced at pH 5.5 (Fig 3H). Concurrently, methionine supplementation not only restored ATP levels but also increased the IC50 of PZA in the *ΔmetZ* at pH 5.5 (Fig 3I and S6F). *M. tuberculosis* produced significantly higher levels of hydrogen sulfide (H_2_S) at acidic pH compared to the neutral pH (Fig S6G). H_2_S serves as a co-substrate and S-donor in the reaction catalyzed by OSHS enzyme (MetZ), an enzyme belonging to the PLP-dependent superfamily of proteins which require PLP as co-factor (Fig 1B and S1). To assess the role of *metZ* in PLP homeostasis, we examined the growth kinetics of different mycobacterial strains in minimal medium supplemented with or without PLP at both pH 7.2 and pH 5.5. We found that knocking out *metZ* had no apparent consequences on the growth kinetics of the *M. tuberculosis* with or without PLP at either pH conditions (Fig. S7A and B). However, we did observe a low pH-based growth defect in *ΔmetZ* even in minimal medium (Fig. S7B). In summary, a defect in the methionine biosynthesis pathway at low pH renders the *ΔmetZ* strain metabolically less active, thereby inhibiting its growth at acidic pH.

### Molecular docking and Molecular dynamics (MD) simulations revealed enhanced stability of MetZ at acidic pH

To investigate the molecular mechanisms underlying MetZ substrate binding and their kinetics at physiologically relevant pH (7.0 and 5.4), we employed computational approaches, including molecular docking and molecular dynamics simulations studies. To perform *in-silico* molecular docking with MetZ protein (Fig. 4A), we determined the binding site of O-succinyl homoserine (OSH) by focused molecular docking [23, 24] that provided the final pose at the cofactor bound site. A pose with docking energy of -4.07 Kcal/mol and cluster size of 50 (out of 200) was found suitable at pH 7.0. While, at pH 5.5 the most likely pose showcased a docking energy of -4.19 Kcal/mol with a cluster size of 80 (200). For both these poses at diverse pH, a static thermodynamic quantification was carried out using MM-GBSA which was found to be -49.20 Kcal/mol and -56.93 Kcal/mol, respectively. Molecular Dynamics Simulation at a constant pH was performed for these poses. Following the 300ns MD simulations, the protein exhibited stable RMSD values at 2.98 Å and 4.34 Å at pH 7.0 and 5.5, respectively. However, significant changes were observed in the stability of the substrate as well as the occupied binding site. On observing the substrate in consensus with its binding site, the region was found to have higher stability and an average RMSD of 1.87 Å was observed at pH 5.5 in comparison to an average of 2.71 Å at pH 7.0 (as shown in Fig. 4B and D). The underlying reason for this higher stability at pH 5.5 is due to the establishment of stable durable bonds in the form of hydrogen bonds with residues V196, F197 and T319 and the O-atoms at the terminals of the substrate. Furthermore, another factor which enhances the strength is the formation of a salt-bridge with residue H220 and substrate. However, a similar configuration of H-bonds and salt-bridge are not observed at pH 7.0, possibly due to conformational dynamics [25] (Fig. 4E). These characteristics are a result of the change in pKa values as well as conformations of the binding site forming amino acid residues. Protonation states of amino acids are highly susceptible to change in environmental pH and thus, preparing the protein at pH 7.0 and 5.5 generates different acid dissociation constants, which ultimately affects the bond forming tendencies of residues. The residue E165 attains a pKa of 8.78 at pH 7.0 and donates a proton, forming a bond with the substrate. Similarly, D194 attains a pKa of 9.11 at pH 5.5 which affects its binding to the substrate. Also, H220 is present in a protonated state at a lower pH and thus involved in the formation of a Salt-Bridge with the substrate. While at pH 7.0, H220 remains in a deprotonated state and thus does not contribute to the stability of the binding pocket at neutral pH. This showcases the importance of the acidic pH which contributes to relatively greater stability of the substrate. On analysis of the binding site, we found that H220 and T319 interact with the substrate at the acidic pH but they showcase conformational changes at pH 7.0 which hinders their interaction. The imidazole ring sidechain of H220 rotates in anti-clockwise direction and thus does not get involved in bond formation. Similarly, T319 forms H-bonds at both pH but the nature of the bonds varies from transient at pH 7.0 to largely consistent at pH 5.5. Overall, conformational changes in the binding site residues can be accredited for better stability at pH 5.5 The stability of the protein and the substrate at acidic pH hints towards a better binding activity of the substrate. This characteristic feature can be examined by performing the MM-GBSA of a few selected frames from both the trajectories to compare their activity. The average binding/biological activity of the protein is found to be -37.35 Kcal/mol and - 59.40 Kcal/mol at pH 7.0 and 5.5, respectively (Fig. 4C). This increase in the binding affinity is due to the effect of acidic pH on the protonation state as well as the induced conformational changes. The docking scores at both pH 7.0 & 5.5 suggest that the residence time of the substrate at the binding site must be short and the sulfur transfer reaction might be transient. Overall, focused docking and MD simulations revealed that although protein is stable at both acidic and neutral pH, the active site of the protein holds substrate (OSH) more stably at acidic pH rather than at neutral pH which sheds light on its physiological role in methionine biosynthesis at acidic pH instead of neutral pH in concordance with our *in-vitro* studies.

**Figure 4.**
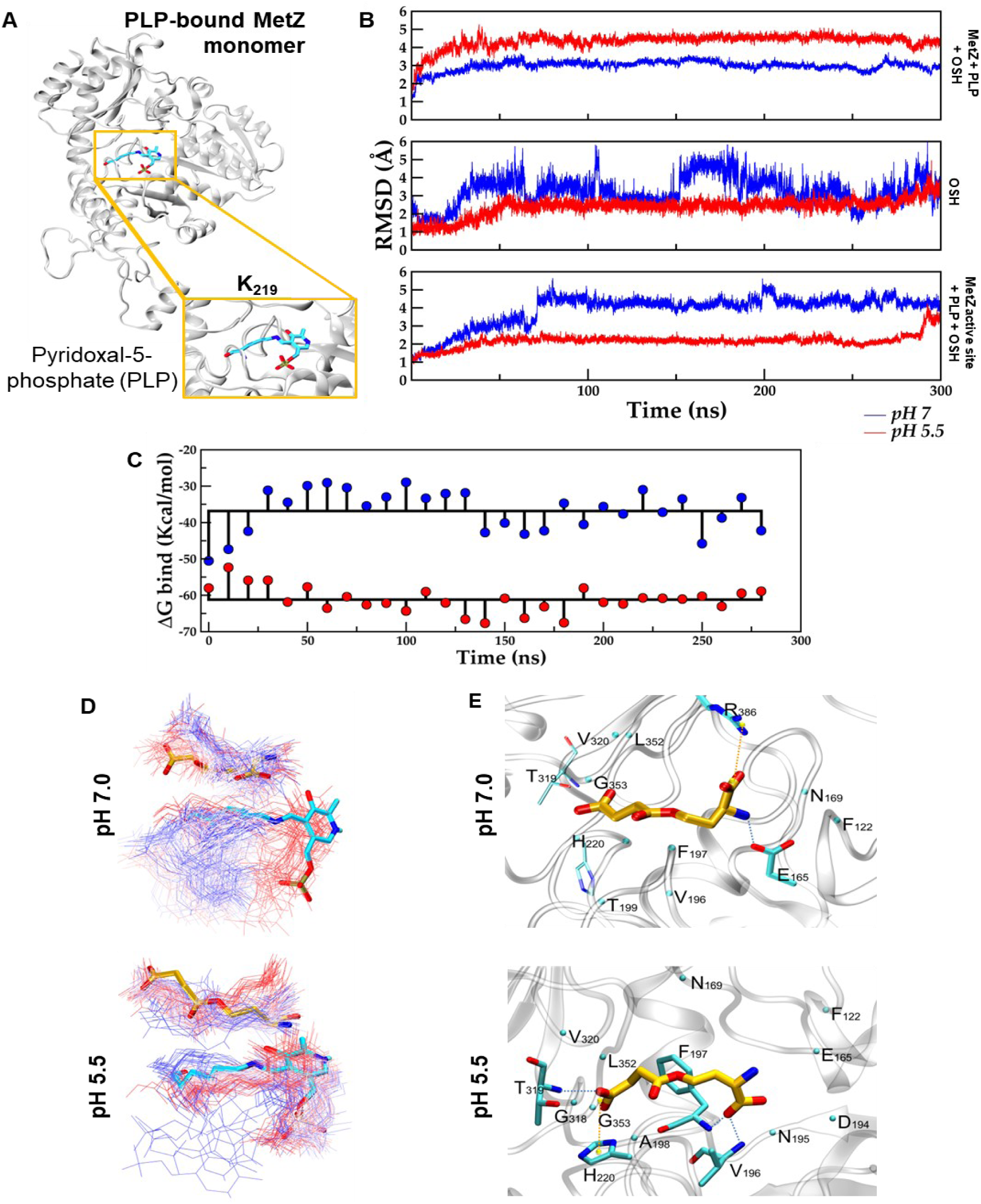
Molecular docking and Molecular dynamics (MD) simulations revealed enhanced stability of MetZ at acidic pH. (A). Structural representation of O-succinyl homoserine sulfhydrylase, rendered in new cartoon and colored in grey (single chain) with pyridoxal-5-phosphate in licorice and rendered atom-wise. (B). Dynamic characteristics of systems elucidated through constant pH MD simulations. The system evolution was measured via RMSD through the course of 300ns of the protein, substrate and the binding site of the substrate. Color coding is followed as the system at pH 7 (blue) and at pH 5.5 (red). (C). The Binding Free Energy (in Kcal/mol) comparison of the complexes at pH 7 (blue) and pH 5.5 (red). (D). The dynamic changes represent the positional shifts of the substrate (orange) and co-factor (cyan) throughout the simulation, with a color gradient (time-dependent manner) from red (initial), and white (intermediate) to blue (final) at pH 7 at pH 5.5. (E). Interaction map of the substrate at pH 7 and pH 5.5. Residues lining the pocket (within 4.0 Å) are shown in licorice and colored atom-wise (C: cyan, N: blue and O: red). The residues forming hydrogen bonds (dotted line in blue) and Salt-bridges (dotted line in yellow) are shown. The Cα atom of residues involved in hydrophobic interaction are shown in beads.

### *metZ*-mediated direct sulfhydrylation pathway is critical for methionine biosynthesis at acidic pH

To investigate the independence of the *metZ*-dependent methionine biosynthesis pathway from the transsulfuration pathway in *M. tuberculosis*, we employed CRISPRi to knockdown *metB* (*Rv1079*) expression in both *Rv-WT* (*Rv::metB_KD_*) and *ΔmetZ (ΔZ::metB_KD_)* strains (Fig. 5A). Anhydrotetracycline (ATc)-induced knockdown of *metB* expression in both strains was confirmed by RT-qPCR analysis (Fig. S8A). While ATc-induced knockdown of *metB* had no significant effect on the growth of the *Rv-WT* strain, it resulted in a growth defect in the *ΔZ::metB_KD_* strain, effectively mimicking a double knockout phenotype (Fig. 5B and S8B). This growth defect is likely attributed to a compromised methionine biosynthesis pathway, as evidenced by a ∼1.5-fold decrease in the intracellular methionine levels in the *ΔZ::metB_KD_* strain compared to the *Rv::metB_KD_* strain (Fig. 5C). Based on these findings we hypothesized that the *ΔZ::metB_KD_* double mutant would exhibit attenuated growth in an animal model due to reduced methionine availability. To test this hypothesis, mice were infected with *Rv::NT* (Non-targeting control sgRNA) and *ΔZ::metB_KD_* strains, and doxycycline was administered to the specific groups to induce CRISPRi-mediated knockdown of the respective gene. Bacterial load was subsequently estimated by CFU plating (Fig 5D). We found that compared to NT and *ΔmetZ*, the *ΔZ::metB_KD_* double mutant exhibited reduced bacterial burden during the initial stages of infection in both lungs and spleen which was partially rescued by week 6 post infection (Fig. 5E). We hypothesized that the presence of an active methionine transporter, such as MetM (Rv3253c) [26] might contribute to the partial rescue of the growth defect observed in the animal infected with *ΔZ::metB_KD_* double mutant strain of *M tuberculosis*. Collectively, the above data demonstrate the functional independence of the MetZ-mediated direct sulfhydrylation pathway in *M. tuberculosis.* The observed growth defects and reduced methionine levels in the *ΔZ::metB_KD_* strain both *in-vitro* and *in-vivo* models highlight the critical role of these redundant pathways in supporting bacterial growth and survival inside the host.

**Figure 5:**
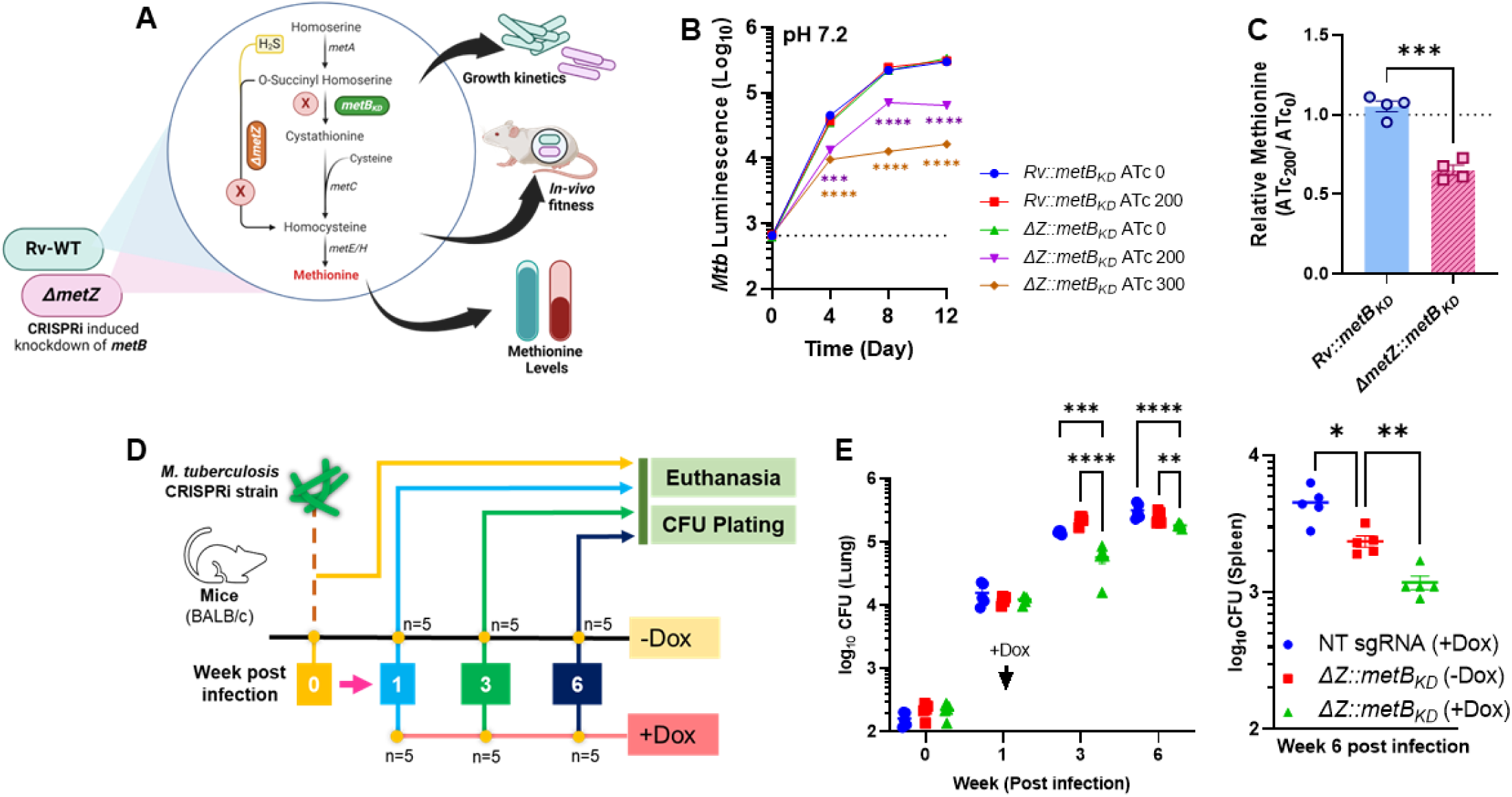
*metZ mediated* direct sulfhydrylation pathway is critical for methionine biosynthesis at low pH. (A). Diagrammatic representation of experimental setup to elucidate independent existence of direct sulfhydrylation pathway. CRISPRi-based knockdown of *metB* (*Rv1079*) strains was achieved in the genetic backgrounds of *Rv-WT* and *ΔmetZ* strains. Upon induction with ATc, these strains were depleted of *metB* transcripts and characterised for *in-vitro* growth kinetics, methionine levels and *in-vivo* fitness. (B). *In-vitro* growth kinetics of CRISPRi knockdown strains. (C). Relative methionine levels in CRISPRi knockdown strains either lacking *metB* (*Rv::metB_KD_*) or both *metZ* and *metB (ΔZ::metB_KD_)* by LC-MS. (D). Graphical representation of *in-vivo* fitness experiment. Mice were infected with CRISPRi knockdown of *metB* in Rv-WT and *ΔmetZ* background and orally fed with doxycycline (1mg/ml) to induce knockdown of *metB* gene in the respective strains. Bacterial burden was determined by CFU plating in lungs and spleen homogenates. (E). and (F). The bacterial burden of mycobacterial strains lacking either *metZ*, *metB,* or both in mice lungs and spleen lysates respectively. Data is represented as mean ± SEM of replicates. Comparisons for statistical significance have been made using ordinary one-way ANOVA (Dunnett’s multiple comparisons test) for (B) where ***P<0.0003, ****P<0.0001; Unpaired t-test for (C) where ***P=0.0001; Mann-Whitney Test for Spleen bacterial burden in (E) where *P=0.015, **P=0.007 and two-way ANOVA (Dunnett’s multiple comparisons) for Lungs bacterial burden in (E) where **P=0.0064, ***P=0.0003 and ****P<0.0001.

### Direct sulfhydrylation pathway of *M. tuberculosis* contributes towards antibiotic persistence in response to Bedaquiline

Next, we wanted to study the role of direct sulfhydrylation pathway in regulating the drug susceptibility profile of *M. tuberculosis*. We first compared the Minimum Inhibitory Concentration (MIC) of various anti-TB drugs. We found that the IC50 values for each of these drugs were similar for all three strains (*Rv-WT*, *ΔmetZ* and the complement strain) (Fig. 6A and 9A-D, F). Further, a time-kill assay was performed to determine the role of *metZ* in modulating the formation of drug-induced persisters in *M. tuberculosis*. The frequency of persisters formation was assessed by comparing the biphasic kill curves generated from the time-kill assays using various antibiotics (RIF, INH, EMB and BDQ) at 10X MIC concentrations. No significant differences in the persisters formation were observed among the strains when treated with RIF, INH or EMB (Fig. 6B). Interestingly, compared to the *Rv-WT* and the complement strains, a significant decrease in the persisters formation was observed in the *ΔmetZ* upon treatment with BDQ at both neutral (Fig. 6B) and acidic pH (Fig. S9E). While a biphasic kill curve was observed for the *ΔmetZ* treated with BDQ at pH 7.2, the kill curve at pH 5.5 was predominantly linear, suggesting a lower frequency of persisters formation at acidic pH. Given BDQ inhibits ATP synthesis, we hypothesized that the observed decrease in the persisters formation in the *ΔmetZ* might be due to impaired generation of ATP. However, ATP depletion kinetics analysis using 2X MIC BDQ revealed higher ATP levels in the *ΔmetZ* compared to the *Rv-WT* and the complement strains (Fig. 6C), suggesting that BDQ-mediated ATP depletion was slower in the *ΔmetZ* strain. In sum, we found that the *metZ*-mediated direct sulfhydrylation pathway modulates the susceptibility of *M. tuberculosis* to BDQ and that the reduction in the BDQ-dependent persisters generation observed in the *M. tuberculosis* strain lacking *metZ* is independent of the ATP levels present inside the bacteria.

**Figure 6:**
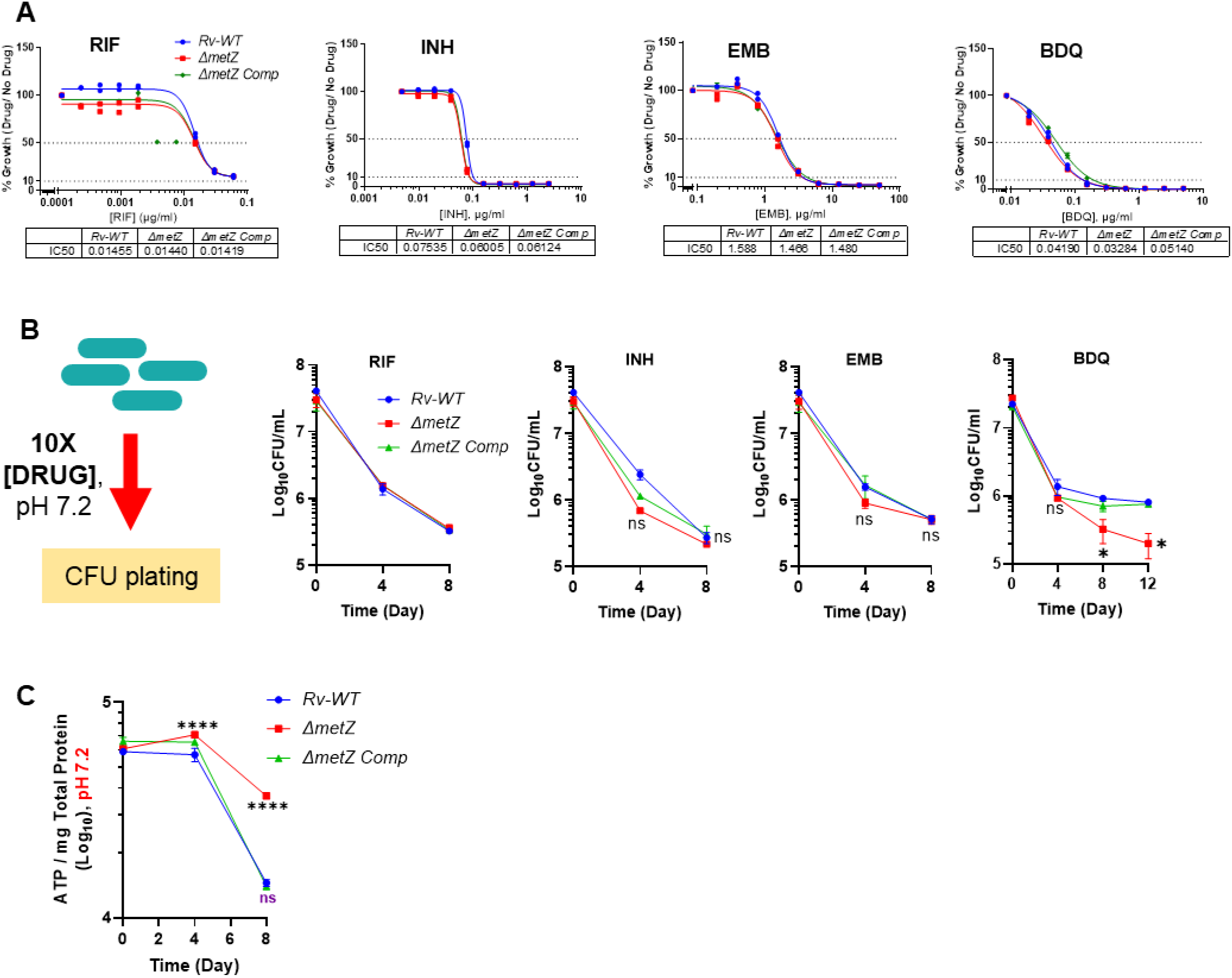
*metZ* is required for persister formation in the presence of BDQ. (A). Minimum Inhibitory Concentration (MIC) determination of the *ΔmetZ* in comparison to Rv-WT and Complement strains against anti-TB drugs: Rifampicin (RIF), Isoniazid (INH), Ethambutol (EMB) and Bedaquilline (BDQ) using microtiter plate assay. IC50 was determined for the drugs with the mentioned strains. (B). Kill kinetics of anti-TB drugs – RIF, INH, EMB, and BDQ at 10X MIC concentrations. Survival was determined in the form of viable bacterial count expressed as CFU. (C). Kinetics of ATP depletion post BDQ treatment. *ΔmetZ* along with Rv-WT and Complement strains were treated with 2X concentration of BDQ and ATP was estimated at indicated time points. *ΔmetZ* maintained high ATP levels. Data is represented as mean ± SEM of replicates. Comparison for statistical significance has been made using two-way ANOVA (Dunnett’s Multiple Comparisons test) for (B) where *P<0.004 and ns = non-significant; two-way ANOVA (Dunnett’s multiple comparisons test) for (B) where *P = 0.029, **P<0.005, ****P<0.0001 and ns, non-significant; using two-way ANOVA (Dunnett’s multiple comparisons test) for (C) where ****P < 0.0001.

### Direct sulfhydrylation pathway supports the growth of *M. tuberculosis* inside the host

To elucidate the role of *metZ* in mycobacterial pathogenesis, a guinea pig model of tuberculosis infection was employed. Briefly, guinea pigs were infected with the respective mycobacterial strains (*Rv-WT*, *ΔmetZ* and complement) via the aerosol route. Bacterial burden in the lungs and the spleen was determined by CFU plating at different time points post-infection. Gross and histopathological analyses were performed to study the damage due to infection at the tissue level (Fig. 7A). We found that at week 4 post-infection the bacterial burden in the lungs of guinea pigs infected with *ΔmetZ* was significantly lower than that observed in animal infected with the *Rv-WT* and complement strains, exhibiting a ∼3.7-fold and 2-fold reduction, respectively. By 8 weeks post-infection, a more pronounced difference was observed, with approximately an 8.9-fold reduction in the bacterial burden in the lungs of *ΔmetZ-*infected animals compared to those infected with the *Rv-WT* strain. Notably, only a modest increase in CFU counts was observed in the *ΔmetZ*-infected animals between weeks 4 and 8 (Fig. 7B). Similarly, a 6-fold reduction in bacterial burden was observed in the spleens of animals infected with the *ΔmetZ* compared to those infected with the WT and complement strains at week 4 post-infection. By week 8, bacterial burden in the spleens of the *ΔmetZ*-infected animals was significantly reduced, approaching the limit of detection by CFU and exhibiting a nearly 263-fold decrease compared to week 4 post-infection (Fig. 7C). Consistent with the reduced bacterial burden (Fig. 7B), gross pathology revealed fewer granulomatous lesions in the lungs of animal infected with the *ΔmetZ* compared to those infected with the *Rv-WT* and complement strains at 8 weeks post-infection (Fig. 7D). To visualize and quantify the damage to lung tissues, Hematoxylin & Eosin (H&E) staining of the formalin preserved lung sections was done. Microscopic examination of lung samples collected from animals infected with *Rv-WT* strain revealed severe granulomatous inflammation throughout the lung parenchyma. Numerous well-defined necrotizing granulomas consisting of epithelioid histiocytes, perimetric cuffing of lymphocytes and plasma cell aggregates and a caseous necrotic mass at the centre were observed. However, lung samples from animals infected with complement strain and *ΔmetZ* strain showed only non-necrotizing granulomas along with multiple foci of inflammatory cell aggregates and occasional alveolar hemorrhagic patches (Fig. 7E). Granuloma scoring further confirmed these observations, with the highest scores observed in lungs sections from animals infected with the *Rv-WT* strain, followed by those infected with the complement and *ΔmetZ* strains (Fig. 7F). Overall, these data suggests that absence of *metZ* gene significantly impairs the ability of *M. tuberculosis* to replicate and survive within the host, suggesting that the *metZ-*dependent direct sulfhydrylation pathway is crucial for the survival of the pathogen inside the host.

**Figure 7:**
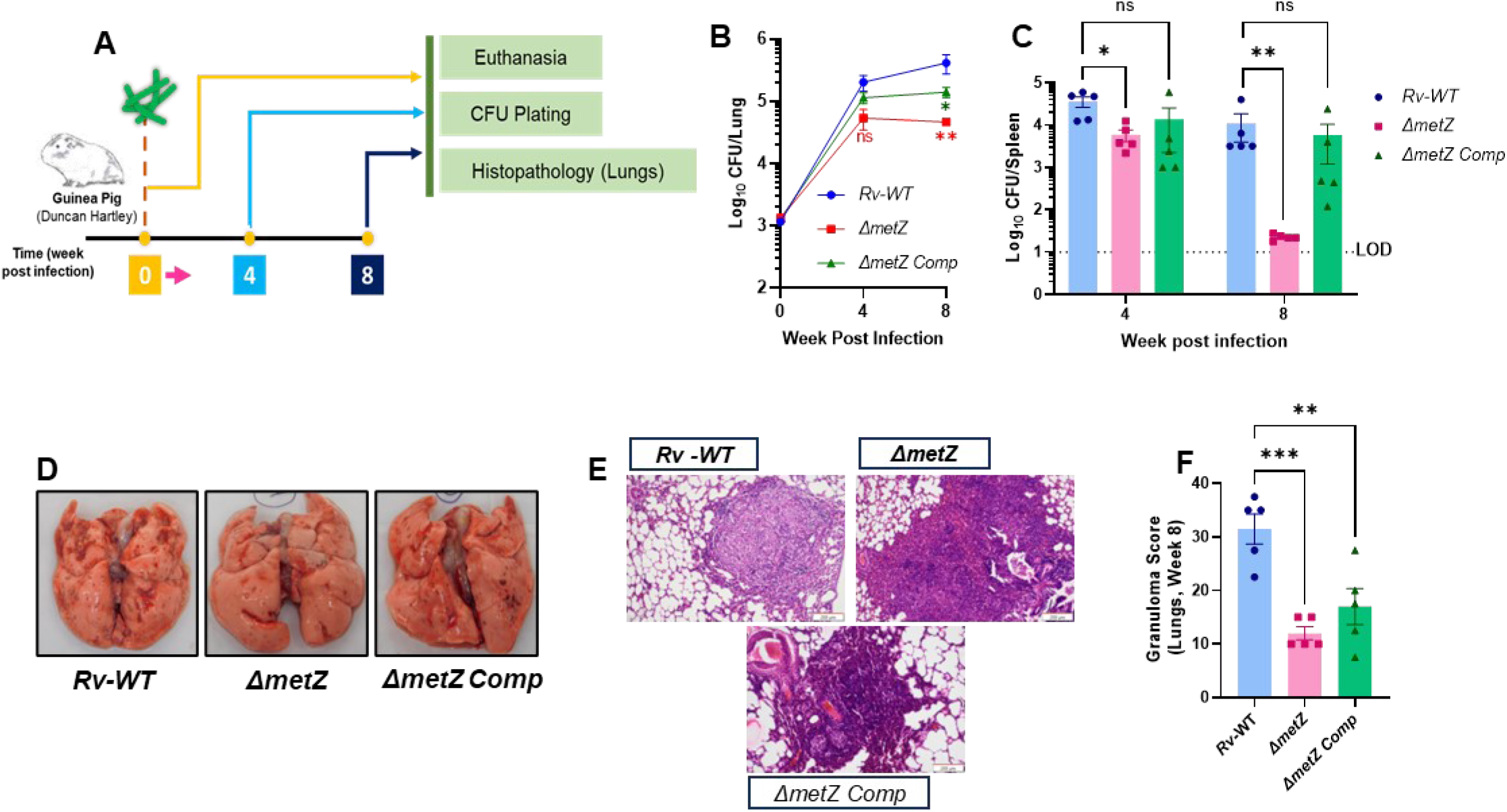
*metZ* is required for survival of *M. tuberculosis in-vivo*. (A). Diagrammatic representation of *in-vivo* fitness experimental set-up. Guinea pigs were infected with WT, mutant, and complement strains, and disease progression was monitored through CFU plating at indicated time points. (B). Bacterial burden in guinea pigs’ lungs as determined by CFU plating of the lung homogenates of the infected animals at indicated time points. The mutant strain failed to replicate in guinea pig lungs post-week 4 of initial infection. (C). CFU burden in spleens of the infected animals revealed that the mutant could not survive inside the spleen post week 4 of infection. (D). Gross lung pathology indicates granulomatous lesions on the infected lungs at week 8-time point post initial infection. (E). H&E staining (F). Total granuloma scores of three strains in lung sections revealed significantly lesser granulomas in *ΔmetZ* infected lungs at week 8 post-infection. Data is represented as mean ± SEM of 5 replicates (animals per group). Comparison for determining statistical significance has been made using the Mann-Whitney Test for (B) and (C) where *P<0.05, **P=0.0079; ordinary one-way ANOVA (Dunnett’s Multiple Comparisons test) for (F) where **P=0.004, ***P=0.0004.

### *metZ* is involved in methionine biosynthesis in *M. tuberculosis* at acidic pH

We hereby summarize that upon infecting macrophages, *M. tuberculosis* traverses a series of intracellular compartments, initially residing in the phagosomes where a relatively neutral pH (∼6.5) and abundant nutrients, including methionine that support bacterial growth. However, as host immunity mounts, IFN-γ-mediated induction of reactive oxygen species (ROS) and reactive nitrogen intermediates (RNI) triggers phagosomal maturation leading to a severe drop in the pH (5.5 or lower). These acidic conditions, coupled with nutrient restrictions imposed by the host, create a hostile environment that inhibits the growth of the pathogen. In response to these stressors, *M. tuberculosis* undergoes substantial transcriptional and metabolic reprogramming. One critical adaptation involves the induction of *metZ* expression, which enables the pathogen to survive these challenging conditions. The MetZ-mediated direct sulfhydrylation pathway utilizes host-derived H_2_S as a co-substrate and S-donor for methionine biosynthesis within this acidic microenvironment, facilitating the maintenance of ATP levels, mycothiol recycling and overall metabolic activity (Fig. 8).

**Figure 8.**
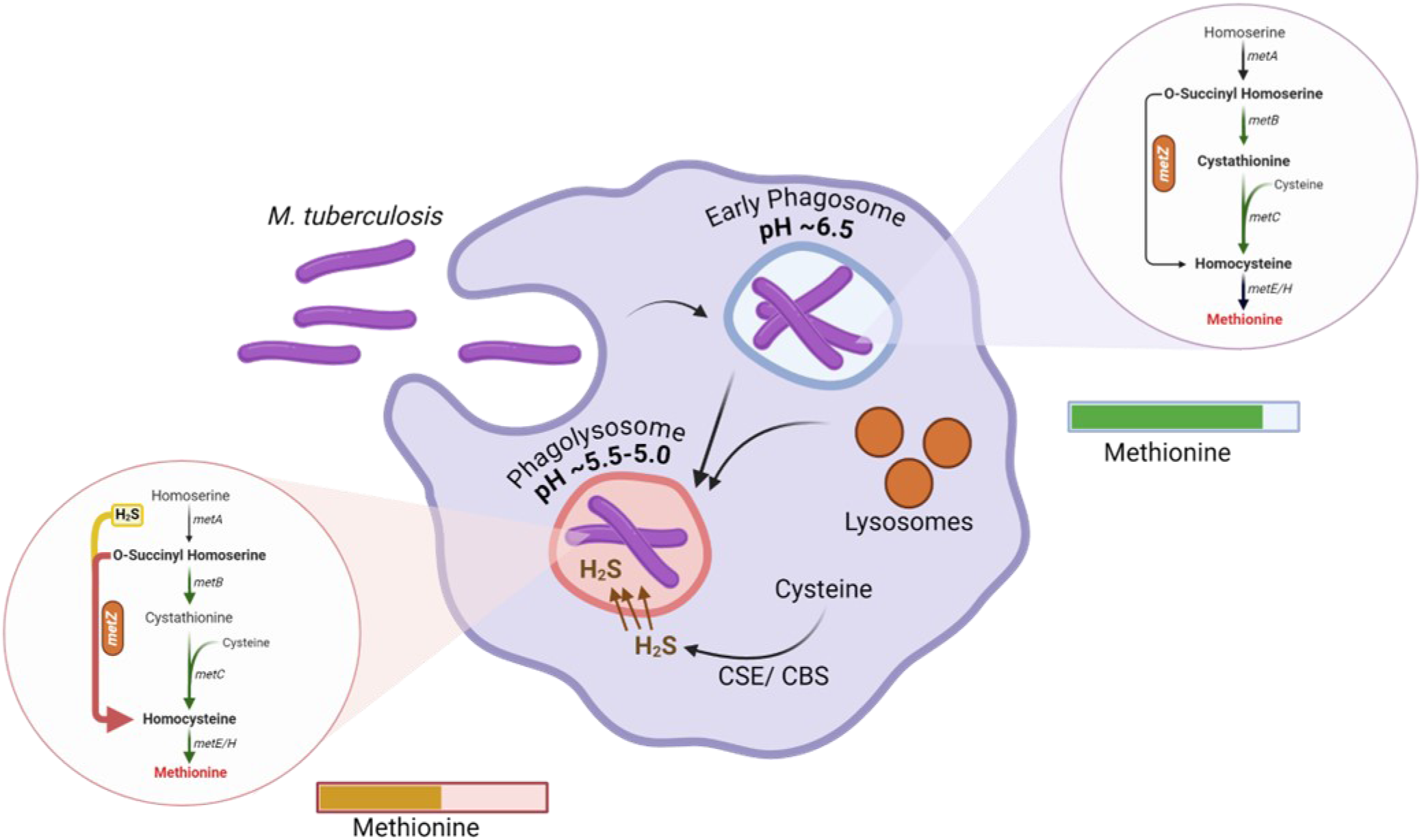
*MetZ-driven* direct sulfhydrylation is critical for methionine biosynthesis at low pH. When *M. tuberculosis* infects macrophage, it passes through a series of compartments starting from the phagosome where pH is essentially ∼6.5 and the bacteria are replicating, with enough nutrients (including methionine). Further, as host immunity kicks in IFN-γ induced production of ROS and RNS followed by phagosome maturation leads to a drop in pH to 5.5 and beyond. Such conditions are unfavourable for *M. tuberculosis* and the host begins to quench nutrients as a means to restrict the trapped bacteria. At this stage bacteria undergo several transcriptional and metabolic changes to sustain the host-induced insults, one of these, as per our findings, includes the induction of *metZ* expression. This helps bacteria survive these stress conditions and maintain methionine levels by utilizing endogenous as well as host-generated H_2_S which acts as co-substrate and S-donor for methionine biosynthesis in acidic micro-environment. This enables bacteria to maintain sufficient levels of methionine to keep adaptation going while also helping the recycling of ATP, mycothiol, at pace with its metabolic activity.

## Discussion

*M. tuberculosis*, adapts to various hostile conditions within host macrophages resulting in the differential expression of proteins crucial for its survival. One such key adaptation involves increase in the biosynthesis of cysteine which compensates for the increase in the flux of cysteine towards generating the thiol-active metabolites (mycothiol, ergothioneine, NAD+, etc.). These metabolites are essential for maintaining the redox balance upon exposure to heightened levels of ROS, RNI [27] and acidic pH [28]. The same has been reported in *E. coli*, where increased oxidative stress leads to higher demand for the cytosolic glutathione which stimulates cysteine biosynthesis [29]. Under such demanding circumstances, the pathogen needs to evolve and devise a mechanism to efficiently regulate the flux of cysteine towards either biosynthesis of methionine and/or mycothiol which would maintain the total thiol pool thereby maintaining redox homeostasis. Likewise, given the crucial role of methionine in bacterial physiology, there has been a gap in our understanding of the rationale of harbouring multiple methionine biosynthesis pathways and their role in maintaining methionine homeostasis critical for the survival of *M. tuberculosis* in a nutrient-deprived *in-vivo* growth environment. We, in the current study, have used a combination of genetics and computational biology-based tools to probe the role of the *metZ-*mediated direct sulfhydrylation pathway in *M. tuberculosis*. We found that MetZ-dependent direct sulfhydrylation is significantly induced at acidic pH. Disrupting this pathway in *M. tuberculosis* results in a decrease, but not complete elimination, of cellular methionine at low pH. Finally, we demonstrated that MetZ-dependent direct sulfhydrylation is required for growth of *M. tuberculosis* inside the host.

In contrast to an earlier report [19], *metZ* was found to be dispensable for the growth of *M. tuberculosis* on cholesterol. The conditional essentiality phenotype observed for *ΔmetZ* under low pH further suggests that the earlier observation of the *metZ* essentiality on cholesterol likely resulted from the low pH growth conditions used in that study. High demand for cysteine, a common precursor for the generation of both methionine and mycothiol, during stress resulted in observed imbalance in the redox homeostasis in *ΔmetZ*. Exogenous supplementation of methionine alleviates the stress by reducing the need to divert the intracellular cysteine pool towards methionine production, thereby enhancing the mycothiol generation critical for mitigating the ROS levels at acidic pH. Beyond redox homeostasis, our data underscores the role of methionine biosynthesis, through the direct sulfhydrylation pathway, in regulating the cytosolic pH and metabolism. The inability of the *ΔmetZ* to maintain the intracellular pH homeostasis impedes the ability of the pathogen to utilize the cellular ATP, leading to a metabolic slowdown. The elevated levels of ATP observed in *ΔmetZ* at acidic pH, is likely, a consequence of reduced ATP utilization rather than overproduction as reported in an earlier study using *E. coli* [30]. The enhanced susceptibility of *ΔmetZ* to Pyrazinamide (PZA), an anti-TB prodrug that gets activated upon exposure to acidic pH, further confirms the presence of a relatively acidic cytosol and accentuates the role of *metZ* in maintaining the pH homeostasis in *M. tuberculosis*. In an *ex-vivo* setting, IFN-γ-induced activation leads to acidification of the pathogen containing phagosomes in macrophages. Consequently, *ΔmetZ* replication inside both resting and the activated macrophages were reduced, likely due to increased co-localization of *ΔmetZ* to acidic compartments as revealed by confocal microscopy. This finding aligns with previous studies suggesting that enhanced susceptibility to oxidative stress results in increased bacterial killing in the acidified macrophage compartments [31-33].

Our data provides a mechanistic understanding of the methionine starvation-associated susceptibility in *M. tuberculosis* in line with reports published by other groups [4, 10]. The presence of both transsulfuration and direct sulfhydrylation pathways for the biosynthesis of methionine in *M tuberculosis* is indeed intriguing. Although both the pathways demonstrate redundancy at normal pH, our qPCR data, along with previous studies, categorically indicate that enzyme responsible for carrying out direct sulfhydrylation is over-expressed in *M tuberculosis* upon induction with acidic pH [32, 34, 35]. These findings are corroborating well with our pH dependent molecular dynamics (MD) simulation studies which suggested that the binding affinity between the substrate and MetZ active site is stronger at low pH. The MD simulation data provide structural insights into how pH and the protonation states of key residues alter the hydrogen bonding and electrostatic interaction network leading to significant changes in the conformation of the MetZ protein. The change in the hydrogen bonding network, identified as a determinant, majorly impacts the change in the binding affinity of the substrate. Furthermore, a comparative analysis of MetZ structural stability at different pH values, using metrics such as root-mean-square deviation (RMSD) and thermodynamic changes, confirms the pH-dependent behaviour of MetZ. The observed changes in the hydrogen bonding network significantly impact the substrate binding affinity of the MetZ, indicating that, under low pH conditions, *metZ*-mediated direct sulfhydrylation majorly contributes towards the biosynthesis of methionine in *M. tuberculosis*. We further confirm that *M. tuberculosis* strain that lacks both transsulfuration and direct sulfhydrylation pathways (*ΔmetZ/metB_KD_*) fails to generate methionine, demonstrating methionine auxotrophy under *in-vitro* conditions. This finding confirms the redundancy that exists between these two pathways. Surprisingly, the growth defect observed in the *ΔZ::metB_KD_* under *in-vivo* conditions, while significant, was less pronounced than that observed *in-vitro*. This may be attributed to the presence of a potential methionine transporter, encoded by Rv3253c (MetM), which allows the pathogen to salvage methionine from the host environment [26]. Similar to our finding (Fig. S6G), it has been reported previously that *M. tuberculosis* can produce H_2_S, a gaseous signalling molecule, upon induction with acidic pH [13] probably as a mechanism to counteract increasing ROS and dropping metabolic potential of the cells. Further, it has also been shown that H_2_S at sub-inhibitory levels in *M. tuberculosis* have a positive effect on the mycobacterial electron transport chain (ETC) and this effect is reversible [36, 37]. Additionally, host-generated H_2_S has been shown to enhance *M. tuberculosis* replication inside the host. This effect is evident in studies demonstrating impaired *M. tuberculosis* replication when the hosts’ ability to generate H_2_S is compromised, either through genetic knockout of the genes encoding the enzyme cystathionine-*γ*-lyase (CSE) and cystathionine-*β*-synthase (CBS) or by using pharmacological inhibitors of these enzymes [36, 38, 39]. Co-incidentally, it is well established in the literature that sulfhydrylases, including MetZ, utilize H_2_S as a S-donor for various reactions, such as cysteine biosynthesis or, in this case of MetZ, for the production of homocysteine. The development of antibiotic persistence in bacteria is a significant contributor to the failure of anti-TB treatment. The frequency of generation of the persisters population can be quantified through a time-kill assay using high antibiotic concentration (5-10X MIC) [40]. Our time-kill experiments suggest that the absence of a direct sulfhydrylation pathway in *ΔmetZ* prevents the BDQ-specific generation of antibiotic persistence in *M. tuberculosis*. Intriguingly, despite exhibiting higher cytosolic ATP levels compared to the wild-type strains, *ΔmetZ* demonstrated a decrease in the BDQ-mediated persisters formation. This observation contrast with a previous study on *ΔpckA* mutant, wherein elevated cytosolic ATP levels increased drug tolerance when grown in acetate medium [41]. In summary, our findings suggest that beyond its role in maintaining methionine levels at acidic pH, the direct sulfhydrylation pathway also modulates the generation of BDQ-specific generation of antibiotic persistence during infection. Further investigation into this complex phenotype will be crucial for developing new strategies to prevent the emergence of BDQ resistance in tuberculosis. Furthermore, we found that *metZ* is essential for the maintenance of *M. tuberculosis* infection in guinea pig model, as evident by a reduced bacterial burden and severe attenuation in the spleen. Spleen is known to be iron rich, which can contribute to the generation of reactive oxygen species via Fenton chemistry. Given the observed susceptibility of the *ΔmetZ* to oxidative stress, the low bacterial count in the spleen at week 8 post-infection likely reflects the inability of the mutant to survive in this nutrient-deprived, low pH and oxidative stress-inducing environment within the host. Our results indicate that *M. tuberculosis* adapts to the decreasing pH in its surroundings by maintaining methionine biosynthesis through direct sulfhydrylation utilizing exogenous and endogenous H_2_S as S-donor.

This study provides a foundation for designing novel anti-tuberculosis treatment strategies. Using pharmacological inhibitors of MetZ as an adjunct to the existing anti-tuberculosis regimens could offer a promising approach. This synergistic strategy can potentially enhance the efficacy of the existing treatment regimen against tuberculosis.

## Materials and Methods

### Bacterial strains

The knockout strain of *metZ* gene *(Rv0391)* was generated in the genetic background of wild type *M. tuberculosis* H37Rv by genetic recombineering. Briefly, a gene-specific allelic exchange substrate (AES) harboring 1kb sequences of target gene upstream and downstream homology flanking Hygromycin resistance gene cassette was constructed in the suicidal vector pJM1. AES was transformed into H37Rv (wild-type, *Rv*-WT) strain overexpressing Phage Che9c RecET proteins to facilitate recombination. Recombinants with the Hygromycin resistance (Hyg^r^) gene were selected over Hygromycin (Hyg) plates and screened using gene-specific PCR. The marked mutant thus prepared was unmarked to excise out Hygromycin resistance cassette using Cre-recombinase expressed from plasmid pCre-SacB-Zeo. The unmarked mutant was confirmed by PCR and RT-qPCR to confirm the loss of gene as well as transcripts of *metZ* gene. The unmarked mutant was then transformed with a plasmid expressing a functional copy of the *metZ* gene under P*_myc1_tetO* promoter replicating either episomally or integrated to generate genetic complementation strain (denoted as *ΔmetZ Comp*). Bioluminescent mycobacterial strains were generated by transforming respective mycobacterial strains (**Table S3, List of strains used in the study**) with integrative plasmid expressing *luxABCDE* operon (Addgene# 26161) [42] with slight modifications [43]. Strains were checked for bioluminescence and a luminescence vs CFU plot was prepared with *Rv-WT* luminescence reporter strain to determine the relation between luminescence and CFU-based read-outs of bacterial viability (Fig. S3A). Henceforth, where ever appropriate, luminescence was used as proxy for the bacterial viability and growth.

### Media and cultural conditions

The bacterial strains were maintained as frozen glycerol stocks at -80°C. Whenever required, the stocks were thawed and diluted 1:500 in 7H9 (Difco, BBL) medium supplemented with Oleic acid, Albumin, Dextrose, Salt (NaCl) (OADS) and 0.01% Tween 80. Bacterial enumeration was done on 7H11 (Difco BBL) agar plates supplemented with OADS. Kanamycin (Sigma), 25μg/ml, Hygromycin (Sigma), 50μg/ml and Zeocin (Invitrogen), 25μg/ml were used whenever required. For media with no carbon source, 7H9 base medium was prepared without glycerol and OADS (or Albumin, Dextrose, Salt: ADS) enrichment and later supplemented with desired Carbon (C)-sources and 0.05% tyloxapol. 100mM stock of Cholesterol (Sigma) was prepared by heating cholesterol in Tyloxapol (Sigma): Ethanol (Merck) mix (1:4) at 80°C for 30 mins [44]. When required, cholesterol was diluted to desired concentration (200μM) in 7H9 base medium wherever specified. Likewise, glycerol (Sigma) was used at a final concentration 0.1% v/v and sodium propionate (Sigma) was used at a concentration of 10mM (0.1%) for growth kinetics studies. For pH-dependent growth kinetics experiments, 7H9 base medium was prepared, supplemented with 0.1% glycerol (v/v) and ADS acidified to desired pH using HCl. For Pyridoxal-5’-Phosphate (PLP) supplementation-based growth kinetics, minimal medium (0.5g/litre asparagine, 1g/litre KH_2_PO_4_, 2.5g/litre Na_2_HPO_4_, 50mg/litre ferric ammonium citrate, 0.5g/litre MgSO_4_.7H_2_O, 0.5mg/litre CaCl_2_, and 0.1mg/litre ZnSO_4_) containing 0.1% glycerol (vol/vol) was used with or without 50μg/ml PLP (Sigma) supplementation. Phosphate buffered saline (PBS) was used at 1X concentration supplemented with tyloxapol (0.05%), PBST whenever mentioned. Bacterial enumeration by CFU was done on 7H11 agar plates supplemented with OADS, wherever mentioned. Colonies were counted after 21 days of incubation. Anhydrotetracycline, ATc (Caymen) was used at 200-300ng/mL concentration unless mentioned otherwise. L-methionine (Sigma) was supplemented at 100μg/ml concentration wherever required.

### CRISPRi knockdown

For transcriptional knockdown of *metB (Rv1079)* using CRISPR interference (CRISPRi), target-specific sgRNA was cloned into *BsmBI* (NEB) digested backbone plJR965 (Addgene # 115163) to generate knockdown construct. The knockdown construct was transformed into *Rv-WT* and *ΔmetZ* strains through electroporation. To achieve knockdown of the target genes, cells were grown to mid-log phase and diluted back to OD 0.01 in fresh culture medium with or without ATc. Growth was monitored by measuring OD of the cultures. Further, expression of target genes was quantified by qPCR to determine the levels of knockdown.

### RT-qPCR

Total RNA was isolated from the respective bacterial strains under specific conditions using Trizol (TaKaRa) according to the manufacturer’s protocol. Briefly, bacterial cells were washed with 1X PBS and resuspended in Trizol. Cells were lysed by bead-beating using 0.22µM sized zirconia beads via 5 cycles with intermittent cooling step of 1 minute. Further, RNA was extracted and precipitated using chloroform-isopropanol treatments. Total RNA thus prepared was treated with Turbo DNaseI (Ambion) to digest away the DNA contamination. 3000 μg equivalent DNase treated RNA was used to prepare cDNA using SuperScript IV Reverse Transcriptase (Invitrogen). Finally, 1μg equivalent cDNA was used to set up qPCR reactions with SYBR (TaKaRa). *sigA* was used as internal control and ΔΔC_t_ method [45] was used determine relative gene expressions w.r.t. specified control conditions. For qPCR at acidic pH, *icl1* was used as positive control.

### Growth kinetics

Auto-bioluminesc ent bacterial strains were grown till OD 0.5 (mid-log phase) and washed once with 1X PBST to remove the spent media. The cells were resuspended in 1X PBST and diluted back to an OD of 0.01 in 7H9 base medium supplemented with different carbon sources – glycerol (0.1%), cholesterol (200µM) or propionate (10mM) for C-source dependent growth kinetics. For pH-dependent growth kinetics, bacterial cells were diluted back to an initial OD of 0.05 in 7H9 medium (supplemented with ADS and 0.2% glycerol) acidified to different pH – 7.2, 6.5, 5.5 and 5.0. Whenever required, L-methionine (100µg/ml, Sigma) was added to the cultures and 50µg/ml L-methionine was replenished every 4 days until the conclusion of the experiment. For CRISPRi based growth kinetics, Anhydrotetracycline chloride (ATc) was added to the cultures (200ng/ml) and replenished every 4 days to maintain the suppression of the target gene until the conclusion of experiment. The growth was monitored by measuring OD and/ or auto-bioluminescence using a Multimode Microplate reader (SpectraMax Pro ID3, Molecular Devices). At times, bacterial growth was also determined by CFU at indicated time points by plating aliquots of cultures and plating appropriate dilutions on 7H11 agar plates.

### Physiological stress survival assay

Bacterial strains were grown till OD 0.5 (mid-log phase) and washed once with 1x PBST. Aliquots of 2ml cell suspensions equivalent to OD 0.2 were treated with 0.5mM H_2_O_2_ (oxidative stress), 200µM Deta-NO (nitrosative stress), 0.01% SDS (membrane stress), PBST (nutrient starvation stress), Phosphate citrate buffer, pH 5.5 (acidic stress) *in-vitro*. The survival of the respective strains was determined by CFU plating at indicated time points and calculating survival index (i.e., CFU of strain/ CFU of *Rv-WT*).

### Drug susceptibility (MIC determination)

Auto-bioluminescent bacterial strains were grown till OD 0.5 (mid-log phase) and washed with 1x PBST. Anti-TB drugs: Rifampicin (RIF), Isoniazid (INH), Bedaquiline (BDQ), Ethambutol (EMB), Clofazimine (CFZ) and Pyrazinamide (PZA) were serially diluted on a 10-points, 2-fold format and added to the 96-well plates (Corning). Cells were added at final density of OD 0.01 to the wells containing drugs and incubated at 37°C for 3 days as described previously [46]. Finally, luminescence was recorded as an indicator of bacterial viability which was expressed as dose-response curves and IC_50_ were calculated using non-linear regression analysis in GraphPad Prism. For MIC determination at pH 5.5, the drug dilutions and the bacterial suspensions (final density of OD 0.05) were prepared in 7H9 medium supplemented ADS (pH adjusted to 5.5). For MIC determination of PZA at pH 5.5 upon supplementation with methionine, an additional group of *ΔmetZ* + Methionine was maintained. Dose-response curves were prepared similarly as described above.

### Kill kinetics

Bacterial strains were grown till OD 0.5 (mid-log phase) and washed with 1x PBST. Aliquots of cell suspensions (2ml) at a bacterial density of OD 0.2 were treated with 10X MIC concentrations of anti-TB drugs-Rifampicin (RIF), Isoniazid (INH), Ethambutol (EMB) and Bedaquilline (BDQ) for mentioned time intervals. Bacterial survival was determined by CFU enumeration at indicated time points. For pH-dependent kill kinetics of BDQ, 7H9 (+ADS) medium acidified to pH 5.5 was employed and 10X MIC BDQ treatment was given as described above followed by CFU plating at indicated time points.

### Methionine estimation

Log-phase cultures of mycobacterial strains were washed with 1X PBST and transferred to 7H9 medium maintained at pH 7.2 or 5.5 at an OD 0.4. Cultures were allowed to grow for 6 days after which they were washed with 1X PBS once followed by washing with chilled methanol. Cells equivalent to OD 1 (∼3x10^8^ cells/mL) were lysed in 1 ml extraction buffer (Acetonitrile:Methanol: water:: 2:2:1) by bead beating with intermittent cooling of 1 min per cycle. Lysates were clarified by spinning at 13,000 rpm, 10 mins at 4°C. The clarified supernatants were filtered through 0.22-µ filters and dried by speed vac to evaporate the extraction buffer. Dried pellets were resuspended in methanol:water (1:1) + 0.1% formic acid and vigorously mixed. Finally, 10μL of the reconstituted samples were run on Hypersil Gold C8 column (100 x 2.1mm) with 0.1% formic acid in water as mobile phase A and 0.1% formic acid in acetonitrile as mobile phase B on SCIEX 6500+ QTRAP and peak and retention time was matched with standard (Methionine) across all the samples. Peak intensities were normalized to CFU of the respective samples.

### Total ATP estimation

Cultures of *Rv-WT*, *ΔmetZ* and complement were grown at acidic pH 5.5 till OD 0.5 (mid-log phase) and washed with 1X PBST. Cells were resuspended in 1000μL 1X PBST (with Protease Inhibitor Cocktail) and lysed by 5 cycles: 45 seconds of bead beating with intermittent cooling on ice. Lysates were passed through 0.5-µ sized filters to remove any bacterial cells or debris. 50μL of clarified lysates were transferred to a 96-well plate (white, flat-bottom, SPL) and mixed (1:1) with BacTiter-Glo (Promega) reaction buffer followed by incubation at 37°C for 30 minutes. 1X PBS was used as blank to cancel out the background signals from the samples. Luminescence was recorded using a Multimode Microplate reader (SpectraMax Pro ID3, Molecular Devices). Normalization was done using total protein content of the respective samples and ATP levels were expressed as relative units w.r.t. *Rv-WT* strain. For ATP estimation of BDQ treated cultures, 0.4 OD bacterial suspensions (2ml) of the respective strains were treated with 2X MIC concentration of BDQ for the indicated time-points and samples were prepared as described above.

For determination of total ATP levels upon supplementation of methionine, the bacterial strains were grown to OD 0.5, washed and transferred to 7H9 (+ADS) pH 5.5 with an additional group of *ΔmetZ* +Methionine. The cultures were allowed to grow for 5 days, washed with PBST and lysates were prepared by bead-beating the cells. ATP estimation was performed as described above.

### Metabolic activity

Auto-bioluminescent mycobacterial strains were grown till OD 0.5 (mid-log phase) and washed with 1X PBST to remove the spent media. Cells were resuspended in 1X PBST and re-diluted to an OD of 0.4 in media acidified to the required pH (pH 5.5). 100μL of these suspensions were added to a 96-well plate (white, flat-bottom, SPL) containing 7H9 (ADS + 0.1% glycerol, pH 5.5) and serially diluted (2-fold). The periphery wells were filled with sterile medium to prevent desiccation. The plate was incubated at 37°C for 5 days after which 20μL of PrestoBlue (Thermo) was added to each well followed by an additional incubation of 2 days to allow the development of color. After 2 days of incubation, luminescence read-out was taken and contents of the wells were transferred to a clear bottom black plate followed by recording fluorescence of AlamarBlue at 560(ex)/ 590(em). Ratio of Fluorescence/Luminescence was calculated to determine metabolic activity per cell.

### Intracellular redox state

*Rv-WT*, *ΔmetZ* and complement strains were transformed with plasmid encoding redox biosensor, Mrx1-roGFP2 as described previously [47]. Cells were grown to OD 0.5 and washed with 1X PSBT and resuspended in PBST in MCTs. *Rv-WT* cells in respective tubes were treated with either 100μM H_2_O_2_ or 100μM DTT as control groups for oxidized and reduced conditions respectively. Further, cells were transferred to 96-well plate (clear bottom, black wells) and excitation at 390 and 490 nm while emission at 510 nm was recorded. Ratio of fluorescence at 390 nm/490 nm were compared to determine intracellular redox state of the cells.

For determination of intracellular ROS levels upon supplementation of methionine, the bacterial strains were grown to OD 0.5, washed and transferred to 7H9 (+ADS) pH 7.2 and pH 5.5 with an additional group of *ΔmetZ* +Methionine. The cultures were allowed to grow for 5 days, washed with PBST and resuspended in a final volume of 1ml PBST at density OD 0.5. The samples were transferred to the 96-well plate (clear bottom, black wells) and excitation at 390 and 490 nm while emission at 510 nm was recorded. Ratio of fluorescence at 390 nm/490 nm were compared to determine intracellular redox state of the cells.

### Lipid hydroperoxidation assay

Total lipids were extracted from the respective bacterial strains as described previously [48]. Briefly, bacterial cells were grown to an OD 0.8-1.0 and pelleted down. Pellets were suspended in methanol:chloroform (2:1), 6 ml overnight at 37°C to extract total cellular lipids. Next day, cells were spun down and supernatants were collected. Supernatants having total cellular lipids were further precipitated by saturating the supernatants with 0.8% KCl. The lower phase was collected and dried in glass vials to evaporate the solvents and concentrate the lipids. Dried lipids were resuspended in 1 ml chloroform (Sigma) and used for FOX2 assay for determining lipid hydroperoxidation in the extracted lipids as described elsewhere [49]. Briefly, 200μL of the concentrated lipids were mixed with 1000μL FOX2 reagent in MCTs in dark for 1hr. Samples were transferred to a 96-well plate and read out at 560 nm.

### Intracellular sulfide estimation

Total sulfide estimation was performed as described previously [50] [37]. Briefly, *Rv-WT*, *ΔmetZ* and complement strains were grown to OD 0.5 (mid-log phase) in regular 7H9 medium supplemented with OADS and 0.01% Tween 80. Cells equivalent to OD 0.5 were inoculated into 7H9 medium supplemented with ADS and 0.01% tyloxapol at pH 7.2 and pH 5.5. After 4 days of incubation, the cultures were pellet down and washed with 1X PBST to remove spent media. Pellets were resuspended in the same media with respective pH 7.2 and 5.5 at an OD 0.4. 100μL of the cell suspensions were added to 96-well plate and mixed with 100μL 2X Bismuth chloride buffer. Following an incubation of 4 hours at 37°C, colorimetry was done at 405nm. The values were normalized to culture OD to determine relative sulfide levels in the respective strains at two different pH.

### Extracellular pH (pH_Ex_) measurement

Extracellular pH (pH_Ex_) of the bacterial cultures was measured using pH indicator dye – Chlorophenol red (CPR) as described previously [28]. Briefly, 7H9 medium supplemented with ADS and 0.01% Tween 80 was used to prepare pH standards – pH 7.0, 6.5, 6.0, 5.5, 5.0, 4.5 and 4.0. To 1ml of these standards, 0.004% v/v CPR was added and colorimetric read-outs at 430nm and 590nm were recorded. Ratio of readouts at 430/590nm were used to prepare a pH calibration curve. For pH of cultures, 250μL of bacterial cultures at indicated time points were taken and spun down at 12,000 rpm for 10 minutes to pellet down bacterial cells. To the supernatants, 0.004% v/v CPR was added and colorimetric assay was performed as described above. Ratio of 430/590nm of cultures was used to extrapolate pH_Ex_ using pH calibration curve. Fresh pH standards were prepared for each time-point.

### Macrophage infection

THP-1 monocytic cells were grown in RPMI medium (Lonza) supplemented with 10% fetal bovine serum (FBS, Invitrogen). Further, cells were seeded at a density of 5x10^5^ cells/ well in a 12-well plate (Corning) and matured for 24 hours with PMA (25ng/mL, Sigma) followed by a rest of 24 hours. Whenever required, cells were activated with IFN-γ (10ng/ml, Peprotech) for 16 hrs before infection. Single cell suspensions of the bacterial strains (*Rv-WT*, *ΔmetZ* and complement) were prepared by soft-spin method as described [43] and used for infecting resting or activated macrophages at an MOI of 1. Bacteria were allowed to infect the cells for 4 hours after which, extracellular bacteria were washed off by three washes with pre-warmed PBS (Gibco). At indicated time points, macrophages were lysed with 1X PBS + 0.01% Triton X-100 to release intracellular bacterial cells. Lysates were serially diluted and plated for bacterial enumeration.

### Confocal Microscopy

THP-1 macrophages were seeded on glass coverslips in a 24-well plate (5x10^5^ cells/well) and were activated by 10ng/ml of human IFN-γ (Peprotech, USA) for 16 hours before infection. Activated THP-1 cells were infected with mycobacterial strains (*Rv-WT*, *ΔmetZ* and complement) harboring Live-Dead reporter plasmid [51] constitutively expressing mEmerld at an MOI of 10. After 4 hours of infection, the cells were washed thrice with 1X PBS to remove the extracellular bacteria and replenished with fresh media. Subsequently, after 24 hours post infection, cells were incubated with 50nM LysoTracker Red DND-99 (Invitogen life Technologies, CA, USA) in complete RPMI media for 30 minutes at 37°C with 5% CO2 and later fixed with 4% paraformaldehyde in 1X PBS. Coverslips were mounted with ProLong Diamond antifade with DAPI (Molecular Probes by Life Technologies, CA, USA). Finally, slides were imaged by Olympus Confocal Laser Scanning Microscope (CLSM). Later, GFP-expressing mycobacteria that colocalized with LysoTracker Red was determined by analyzing the ∼ 100 phagosomes and co-localization index was determined.

### Animal Infection

*In-vivo* experiment in this study was performed with approval from the Institutional Animal Ethics Committee (IAEC Approval No: IAEC/THSTI/165) as per guidelines laid by the Committee for the Purpose of Control of Experiments on Animals (CPCEA), Government of India. Briefly, bacterial suspensions of respective strains at a density of 10^8^ cells/mL were prepared in normal saline and guinea pigs (Duncan Hartley) were infected via aerosol route using Inhalation Exposure System (Glas-Col). To determine baseline infection in all groups, a few representative animals (n=3 per group) were sacrificed, followed by an evaluation of disease progression at indicated time points with animals (n=5) per group; organs were harvested and bacterial burdens were determined by CFU plating. Histopathological damage to the lung tissue was determined by H&E staining of lungs fixed in 5% paraformaldehyde followed by granuloma scoring of the infected lung tissue.

### CRISPRi-mediated *in-vivo* fitness

The experiment was performed with approval from the Institutional Animal Ethics Committee (IAEC Approval No: IAEC/THSTI/196) as per guidelines laid by the Committee for the Purpose of Control of Experiments on Animals (CPCEA), Government of India. Briefly, bacterial suspensions of respective strains (***Rv***::NT and *ΔZ::metB_KD_*) at a density of 3x10^8^ cells/mL were prepared in normal saline and BALB/c mice were infected via aerosol route using Inhalation Exposure System (Glas-Col). To induce CRISPRi-mediated gene silencing of *metB* in *ΔZ::metB_KD_* strain, doxycycline (1mg/ml) was administered to mice via oral route at indicated time points [52]. NT (non-targeting CRISPRi strain) was maintained with doxycycline as control which is isogenic to *Rv-WT*. Animals were euthanized at indicated time points, organs (lungs and spleens) were harvested and bacterial burden were determined by CFU plating.

### Bioinformatics analysis

MetZ protein sequence was retrieved from Mycobrowser (https://mycobrowser.epfl.ch/genes/Rv0391) and the cluster of orthologous groups analysis (COG) was performed using Eggnog v5 [53]. Further, a multiple sequence alignment of the protein sequence from members of *Mycobacterium* genus (*M. tuberculosis, M. bovis, M. canettii, M. marinum, M. leprae* and *M. smegmatis*) along with gram-negative (*Pseudomonas aeruginosa*) and gram-positive bacterium (*Corynebacterium glutamicum* and *Rhodococcus*) was performed using Clustal W [54] to identify common domains and motifs amongst the protein sequences.

### Molecular docking and molecular dynamics (MD) simulations

For system preparation, the crystal structure of putative O-succinylhomoserine sulfhydrylase of *M. tuberculosis* was retrieved from RCSB PDB [55] (PDB-ID: 3NDN [56]). The protein has 4 chains and exists in a homo-tetramer form. The receptor has a covalently bound pyridoxal-5-phosphate attached to a modified lysine (K219). The substrate, O-succinyl homoserine, was retrieved from PubChem (PubChemCID 439406). The substrate undergoes a sulfur transfer reaction in the presence of pyridoxal-5-phosphate forming L-homocysteine which takes part in the methylation cycle. The receptor protein was optimized and then minimized using the OPLS3 force field of Protein Preparation Wizard module [57] of the Schrodinger suite. Next, Binding Site Identification was done using AutoDock 4 [58] wherein blind docking of the substrate in the presence and absence of the cofactor to identify the possible binding site was performed. Autogrid4 was used to build a 126x126x126 points grid map with a spacing of 0.586 angstrom and positioned at the center of mass of the receptor protein. The Lamarckian genetic algorithm was employed to generate 200 runs. On docking the substrate in the presence of the cofactor, no suitable results were produced. However, docking in the absence of the co-factor resulted in comparatively higher binding affinity and a larger cluster size. Focused docking on the identified site was also carried out by generating a grid map of 40x40x40 units with a spacing of 0.375 A and 17X20X22 as grid center coordinates. MM-GBSA of few conformers based on their binding affinity and cluster size was done using Prime module [59] of the maestro. Finally, the lowest energy conformations of the substrate were subsequently used for constant pH MD. Further, Constant pH Molecular Dynamics Simulation (MDS) was performed with the Desmond [60] module from Schrodinger suite. Both systems were protonated at pH 5.5 and 7.0, respectively, and minimized with OPLS-all Atom force fields and solvated with the predefined TIP3P water solvent model. The complexes were placed in the orthorhombic periodic boundary conditions to specify the shape and size of the repeating unit buffered at 12 A distances. 4 Na+ and 9 Na+ ions were added to both the systems respectively, to neutralize the systems. The NPT ensemble was employed for the simulation with a timestep of 0.002 ps. The simulation was carried out for a total of 300 ns for each system and the coordinates were saved at intervals of 20 ps to generate 15000 frames. Also, the protein was simulated without the substrate at the two pH values. The post-processing of MD simulation trajectories was carried out using VMD [61]. The root-mean-square-deviation and fluctuation were measured for the protein, substrate and cofactor. Lastly, for thermodynamics analysis of the system we performed Molecular Mechanics Generalized Born Surface Area (MM-GBSA) analysis to evaluate the MD trajectories using Schrodinger software. To effectively encompass the thermodynamic properties, every 500th frame was selected for MM-GBSA analysis.

## Statistics and Reproducibility

The data values are represented as mean ± SEM of the replicates from two-three independent experiments. Wherever applicable, the number of replicates per group have been indicated by the dots. Statistical comparisons have been made with *Rv-WT* in most cases unless mentioned otherwise. GraphPad Prism (ver 9.0) has been used for statistical analysis of all data with suitable statistical tests mentioned with figure legend.

## Supporting information

List of Plasmids

List of Strains

List of Primers

## Acknowledgements

The authors acknowledge THSTI for infrastructural support, Department of Biotechnology, Government of India for funding, the Experimental Animal Facility (and staff) for supporting animal experimentations and BSL-3 facility (and staff) for conducting all *M. tuberculosis* related experiments. Technical support of Mr. Surjit Yadav is duly acknowledged. We acknowledge the Multi-Omics facility for LC-MS based experiments. We also duly acknowledge the receipt of plasmids and constructs used in the study (as listed in Table S1).

## Supplementary material

1. Table S1: List of plasmids and constructs used in the study
2. Table S2: List of bacterial strains used in the study
3. Table S3: List of primers used in the study
4. Supplementary data

