## Supplementary Data for "pH-dependent direct sulfhydrylation pathway is required for the pathogenesis of *Mycobacterium tuberculosis*"

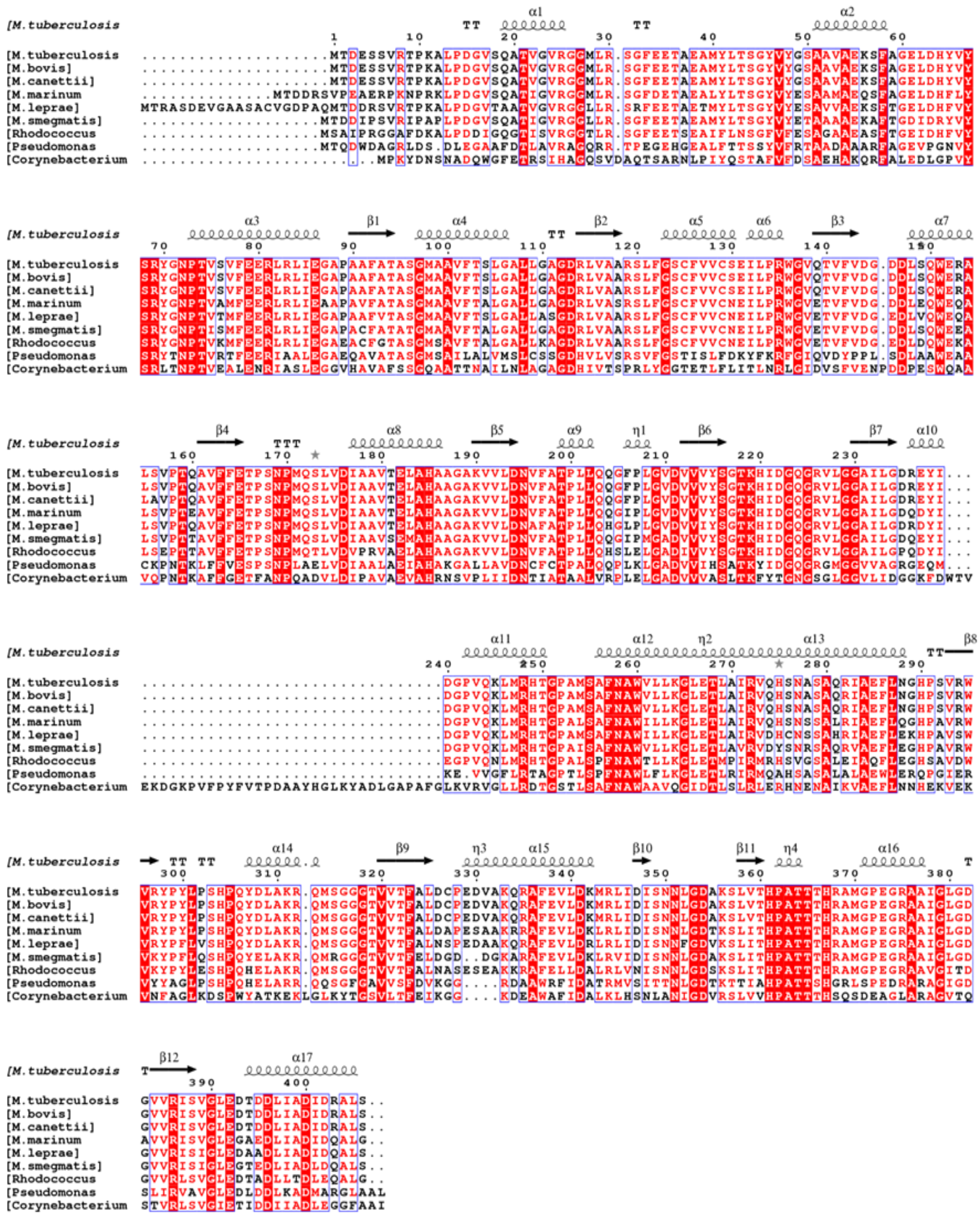

**Figure S1: Rv0391 (MetZ) sequence is conserved across the *Mycobacterium* genus.** (A). Protein sequence alignment of MetZ across other members of MTBC. Multiple sequence alignment (MSA) was done using bacterial species with gram-negative (*Pseudomonas aeruginosa*) and gram-positive bacterium (*Corynebacterium glutamicum* and *Rhodococcus*) along with other mycobacterial species (*M. tuberculosis, M. bovis, M. canettii, M. marinum, M. leprae* and *M. smegmatis*). MSA revealed similarity amongst analysed sequences wrt. Several motifs give rise to a similar secondary structure (indicated by coils on top of the alignment) and conserved lysine (K219) at which the co-factor pyridoxal-5'-phosphate is covalently attached. The presence of a characterized OSHS protein in *C. glutamicum*, in particular with similar motifs to Rv0391 indicates towards the probable role of MetZ (Rv0391) as an OSHS.

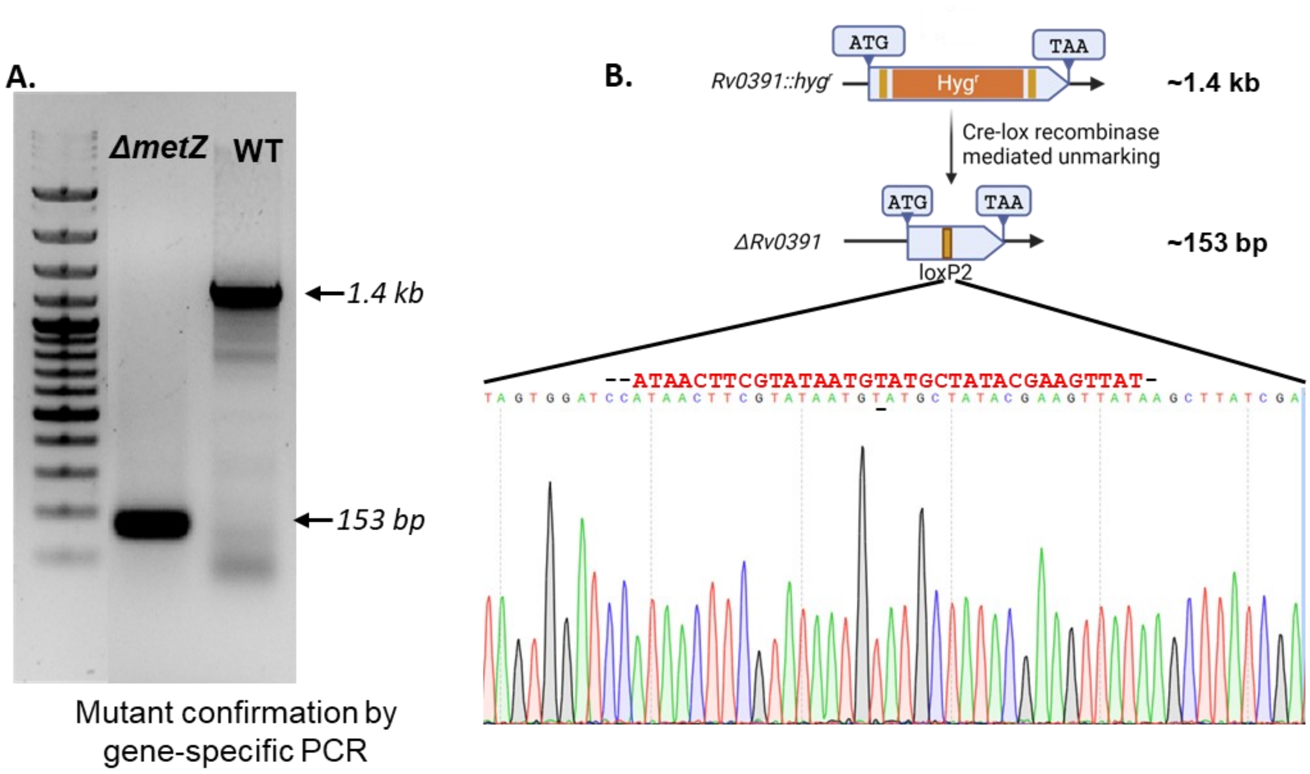


**Figure S2: *Rv0391 (metZ)* genetic mutant confirmation.** (A). Mutant confirmation by gene-specific PCR. Genomic DNA was isolated from the unmarked mutant culture and WT *M. tuberculosis (Rv-WT)* using which a 1.4kb sized fragment encoding *metZ* gene was amplified. The unmarked mutant carried a deletion of a major fragment of the said ORF thus, showing an amplicon of 153bp, confirming the loss of *metZ* encoding sequence. (B). Genetic locus encoding *metZ* was sequenced in the unmarked mutant wherein, loss of the functional ORF confirmed the generation of the mutant when compared to Rv-WT sequence. Post unmarking, a 34bp *loxP* sequence remains in the bacterial chromosome within the target sequence which serves as an indicator of gene deletion and unmarking of the mutant, thereby confirming the loss of the $Hyg^r$ cassette.

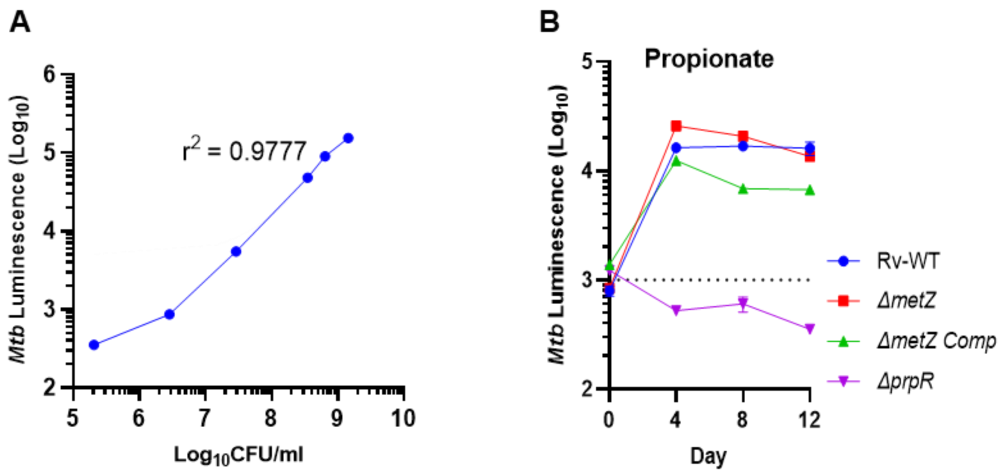


**Figure S3: Generation of auto-bioluminescent *Rv-WT*, *ΔmetZ* and complement stains and growth kinetics.** (A). Linear regression between Luminescence and CFU in *M. tuberculosis*. Log-phase culture of *M. tuberculosis* luminescence reporter strain was grown and ~10 OD equivalent cells were serially diluted (1:10) through 0.0001 OD. For each dilution, bacterial enumeration was done by CFU plating and luminescence was recorded. A linear relationship between luminescence and CFU was determined using linear regression analysis categorically indicating that luminescence can be used as a proxy for bacterial viability and an alternative to CFU. (B) *In-vitro* growth kinetics of *ΔmetZ* in propionate (10mM) as sole C-source. Propionate is a short, odd-chain fatty acid and an end product of cholesterol and fatty acid catabolism in *M. tuberculosis*.

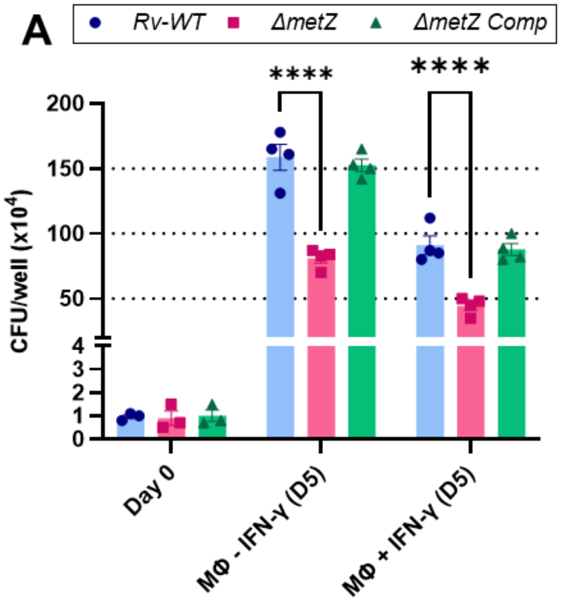


**Figure S4: Intracellular survival assay.** (A). Resting or IFN-γ activated THP-1 macrophages were infected with Rv-WT, *ΔmetZ* and Complement strains and bacterial survival was determined by plating macrophage lysates at the indicated time points. Dots represent the replicates per group and data is shown as mean ± SEM of the replicates. Comparisons have been made with *Rv-WT* for statistical significance using two-way ANOVA (Dunnett's Multiple Comparisons test) where ****P <0.0001. The absence of statistical significance means non-significant changes w.r.t. control (*Rv-WT*). The survival Index calculated from the CFU values is shown in Figure 2D.

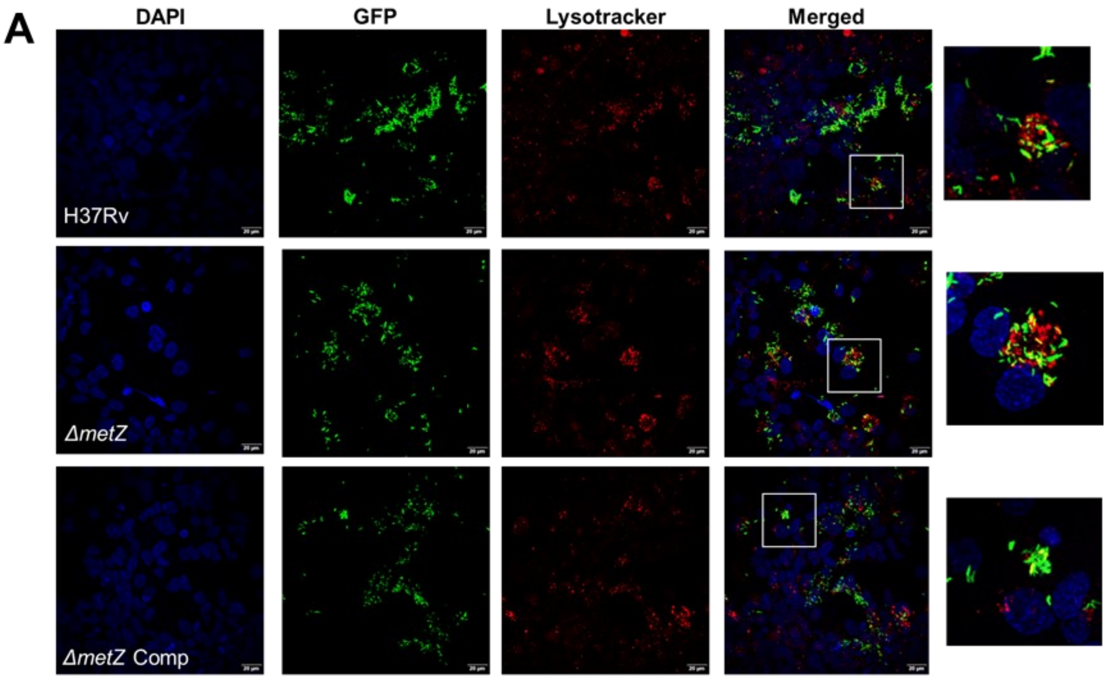


**Figure S5: Panel showing intracellular co-localization of the *Rv-WT*, *ΔmetZ* and Complement strains.** mEmerald-expressing mycobacterial strains were used to infect IFN-γ activated THP-1 macrophages. Two days post-infection, the cells were fixed and stained with DAPI (for the nucleus) and LysoTracker Red. *ΔmetZ* cells were co-localized with acidic regions of the cells (stained in red) as also indicated by a higher co-localization coefficient as shown in Figure 2E. White bars indicate the scale corresponding to 20μM length.

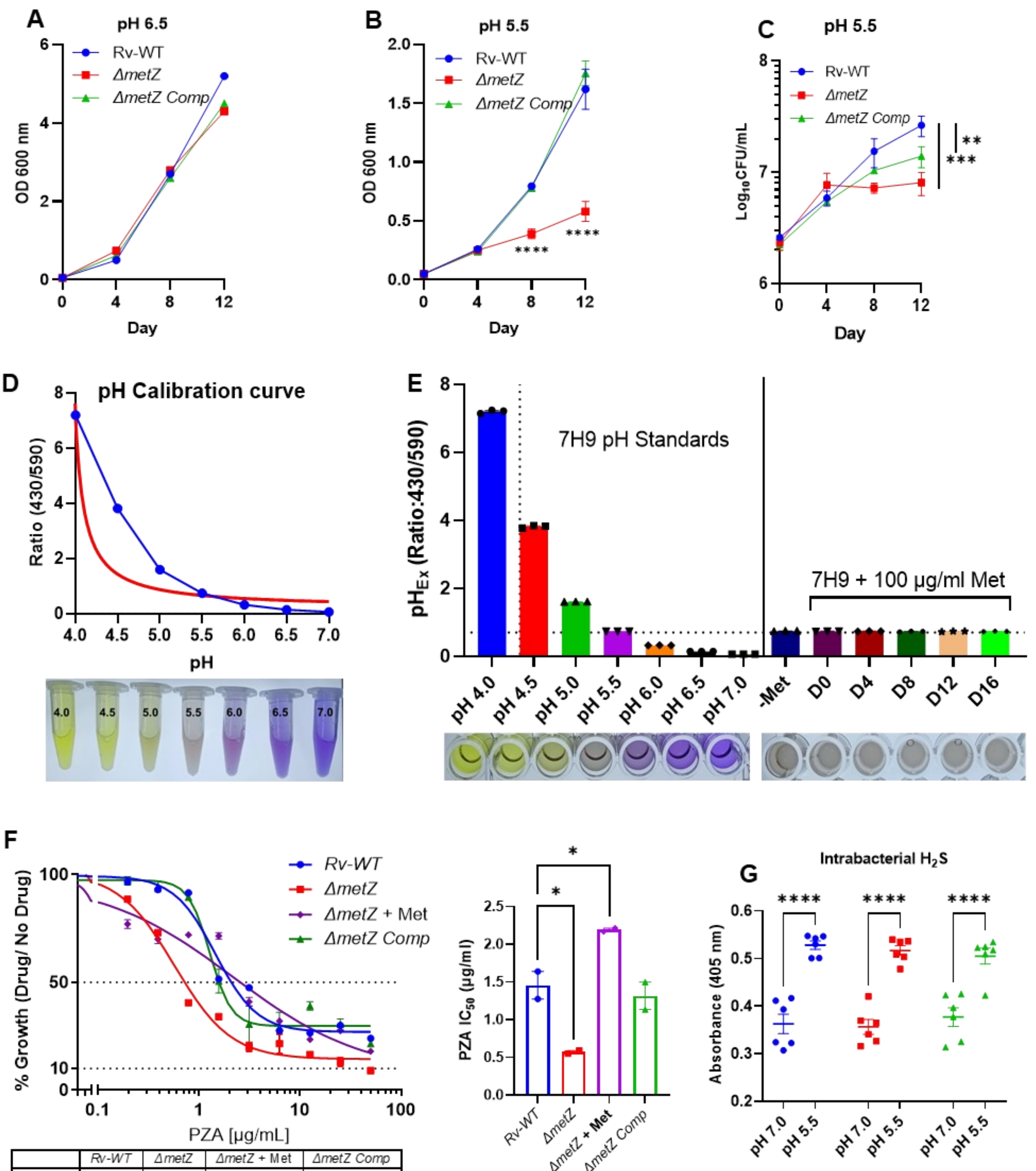


| | Rv-WT | ΔmetZ | ΔmetZ + Met | ΔmetZ Comp |
|---|---|---|---|---|
| IC50 | 1.449 | 0.5687 | 2.183 | 1.276 |

**Figure S6: pH-dependent growth kinetics and methionine supplementation.** (A). *In-vitro* growth kinetics of *ΔmetZ* at pH 6.5. (B). *In-vitro* growth kinetics of *ΔmetZ* at pH 5.5. (C). The viable bacterial count of *ΔmetZ* compared to *Rv-WT* and Complement strains at pH 5.5. (D). pH Calibration curve of 7H9 medium acidified to indicated pHs. (E). Extracellular pH ($pH_{Ex}$) of bacterial culture at various time points compared to pH standards. (F). PZA sensitivity assay of *ΔmetZ* at pH 5.5 with methionine supplementation. IC50 of PZA for the respective strains at acidic pH 5.5 was determined and represented as a dose-response curve prepared by non-linear regression analysis using GraphPad Prism. (G). Endogenous $H_2S$ estimation in bacterial cultures of respective strains at pH 7.2 and 5.5 using Bismuth chloride-based colorimetry assay. Data is represented as mean ± SEM of the replicates. Comparisons for statistical significance have been made with *Rv-WT* using two-way ANOVA (Dunnett's multiple comparisons test) for (A), (B) and (C) where **P = 0.0059, ***P = 0.0003, ****P<0.0001; using ordinary one-way ANOVA (Dunnett's multiple comparisons test) for (F) where, *P=0.01, absence of statistical significance indicates non-significant changes and two-way ANOVA (Sidak's multiple comparisons Test) for (G) were ****P<0.0001.

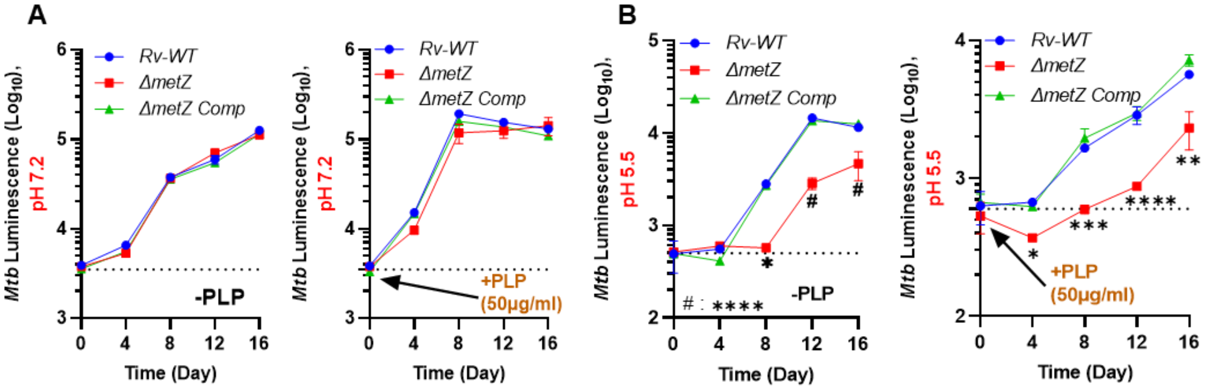


**Figure S7: Effect of *metZ* deletion on PLP homeostasis in *M. tuberculosis*.** (A). *In-vitro* growth kinetics of *ΔmetZ* at pH 7.2 with or without Pyridoxal-5'-phosphate (PLP) supplementation. (B). *In-vitro* growth kinetics of *ΔmetZ* at pH 5.5 with or without Pyridoxal-5'-phosphate (PLP) supplementation. Data is represented as mean ± SEM of the replicates. Comparisons for statistical significance have been made with *Rv-WT* using two-way ANOVA (Dunnett's multiple comparisons test) for (A) and (B) where *P<0.04, **P<0.004, ***P<0.0002, # = ****P<0.0001. No statistical significance indicates non-significant differences.

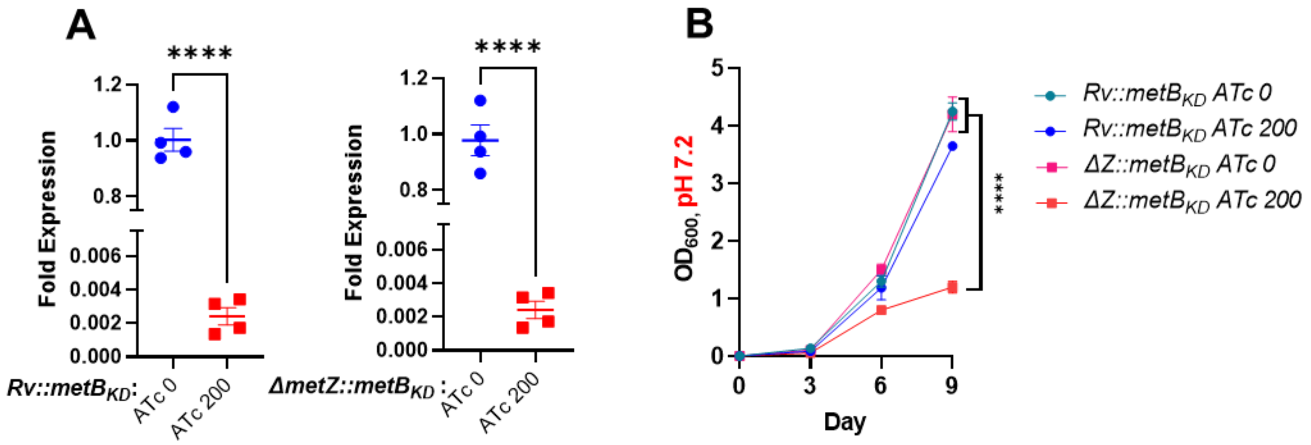


**Figure S8: *metB* knockdown generation and confirmation.** *metB* (*Rv1079*) CRISPRi knockdown construct was transformed into *Rv-WT* and *ΔmetZ* (luminescence reporter strains) to investigate the effect of loss of both transsulfuration and direct sulfhydrylation on bacterial growth. (A). Confirmation of $metB_{KD}$ using RT-qPCR. CRISRPi strains were induced with ATc (200ng/ml) and, fold expression of *metB* w.r.t. ATc untreated cultures of $Rv::metB_{KD}$ and $\Delta Z::metB_{KD}$ strains was determined using the ΔΔCt method. (B). Growth kinetics of *metB* knockdown (OD-based) in $Rv::metB_{KD}$ and $\Delta Z::metB_{KD}$ strains at pH 7.2. Data is represented as mean ± SEM. Comparison for statistical significance has been made using Unpaired t-test for (A), where ****P<0.0001 and two-way ANOVA (Dunnett's multiple comparisons test) for (B), where, ****P<0.0001

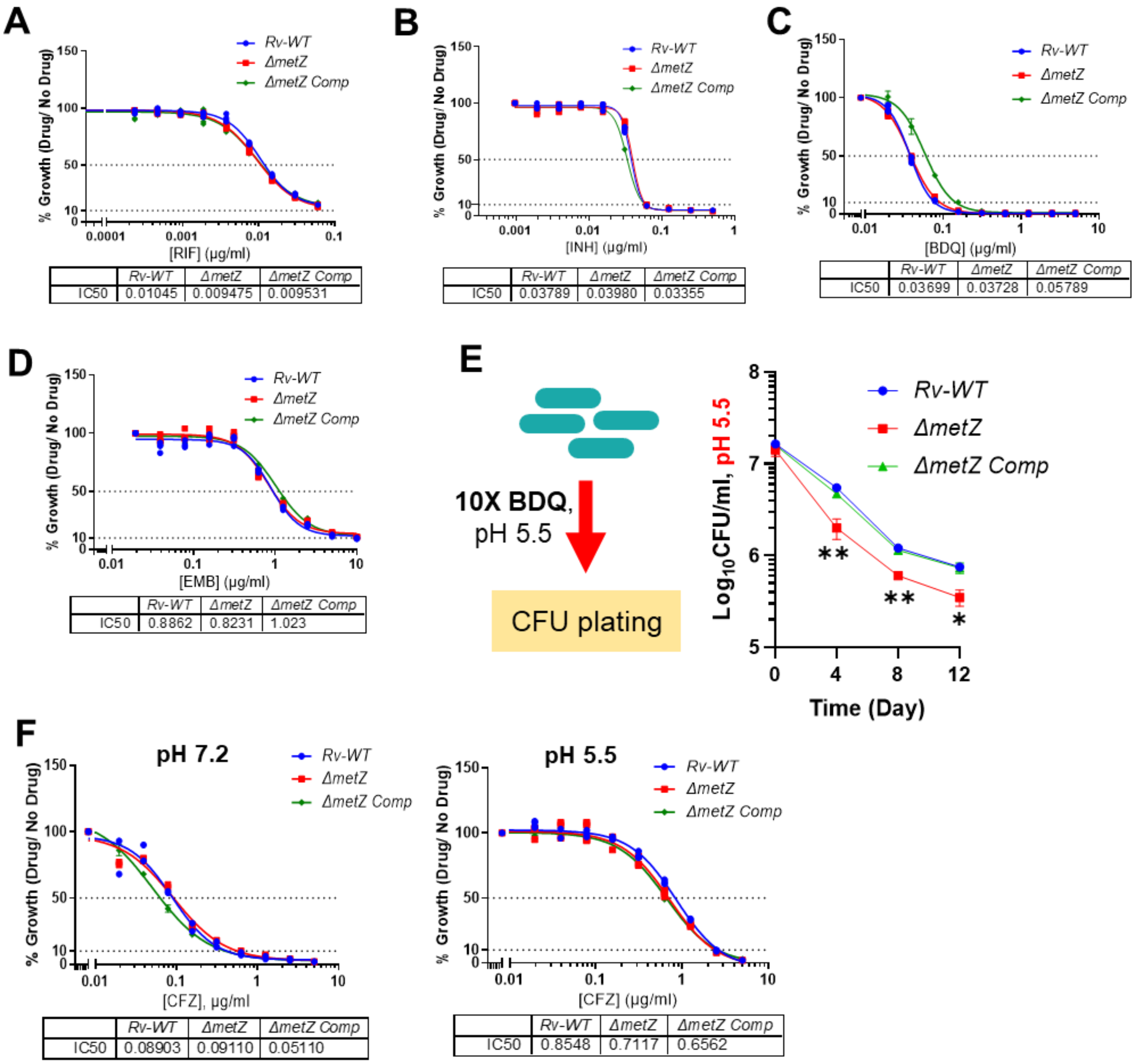


| | Rv-WT | ΔmetZ | ΔmetZ Comp |
|---|---|---|---|
| IC50 | 0.01045 | 0.009475 | 0.009531 |

| | Rv-WT | ΔmetZ | ΔmetZ Comp |
|---|---|---|---|
| IC50 | 0.03789 | 0.03980 | 0.03355 |

| | Rv-WT | ΔmetZ | ΔmetZ Comp |
|---|---|---|---|
| IC50 | 0.03699 | 0.03728 | 0.05789 |

| | Rv-WT | ΔmetZ | ΔmetZ Comp |
|---|---|---|---|
| IC50 | 0.8862 | 0.8231 | 1.023 |

| | Rv-WT | ΔmetZ | ΔmetZ Comp |
|---|---|---|---|
| IC50 | 0.08903 | 0.09110 | 0.05110 |

| | Rv-WT | ΔmetZ | ΔmetZ Comp |
|---|---|---|---|
| IC50 | 0.8548 | 0.7117 | 0.6562 |

**Figure S9: Drug Susceptibility Testing (DST) of anti-TB drugs at acidic pH.** Dose-response curves of Minimum Inhibitory Concentration (MIC) assay at pH 5.5 for anti-TB drugs with IC50: (A). Rifampicin (RIF), (B). Isoniazid (INH), (C). Bedaquilline (BDQ) and (D). Ethambutol (EMB). IC50 values were determined from respective dose-response curves made by non-linear regression analysis using GraphPad Prism. (E). Kill kinetics of BDQ at acidic pH 5.5. (F). MIC assay of Clofazimine (CFZ) at pH 7.2 and 5.5. Data is represented as mean ± SEM. Comparison for statistical significance has been made using Ordinary one-way ANOVA (Dunnett's multiple comparisons test) for (E), where, *P = 0.0249 and **P < 0.005.
