## Supplementary material for "pH-dependent direct sulfhydrylation pathway is required for the pathogenesis of *Mycobacterium tuberculosis*": List of Plasmids

**List of plasmids used in the study**

| S. No. | Plasmid | Feature | Reference |
|---|---|---|---|
| 1. | pJM1 | Hyg/Cm, suicidal vector for generating knockout construct for gene deletion in *M. tuberculosis* | Kind gift from Prof. Christopher M. Sassetti |
| 2. | pST-K | Kan, episomal, mycobacterial expression vector | Kind gift from Dr. Vinay K. Nandicoori, NII, India |
| 3. | pST-Ki | Kan, *attB* integrative, mycobacterial expression vector | Kind gift from Dr. Vinay K. Nandicoori, NII, India |
| 4. | pST-K:*metZ* | Kan, pST-K backbone with *metZ* encoding ORF | This study |
| 5. | pST-Ki:*metZ* | Kan, pST-Ki backbone with *metZ* encoding ORF | This study |
| 6. | pMV261: mEmerald, TagRFP | Hyg, Mycobacterial Live-Dead reporter with ATc inducible RFP | Kind gift from Prof. Christopher M. Sassetti |
| 7. | pJW276 | Zeo, Integrative, Lux expressing plasmid | Kind gift from Prof. Sarah M. Fortune |
| 8. | pMV306:*luxAB*+G13 | Kan, Integrative, Lux expressing plasmid | Addgene # 26161 |
| 9. | pMV762: Mrx1-roGFP2 | Hyg, episomal, genetically encoded redox biosensor | Kind gift from Dr. Amit Singh, IISc, India |
| 10. | plJR965 | Kan, *attB* integrative, target sgRNA and $dCas9_{Sth}$ expressing plasmid for transcriptional knockdown of target gene(s) in *M. tuberculosis* | Kind gift from Prof. Sarah M. Fortune |
| 11. | plJR965::*metB*_sgRNA | CRISPRi knockdown construct for targeting *metB* (*Rv1079*) gene in *M. tuberculosis* | This study |
