## Supplementary material for "pH-dependent direct sulfhydrylation pathway is required for the pathogenesis of *Mycobacterium tuberculosis*": List of Strains

**List of bacterial strains used in the study**

| S. No. | Strain | Marker | Feature | Reference |
|---|---|---|---|---|
| 1. | *E. coli* XL1Blue | | | This lab |
| 2. | *M. tuberculosis* H37Rv | | Wild type *Mtb (Rv-WT)* | Kind gift from Christopher M. Sassetti |
| 3. | *ΔprpR::Hyg$^r$* | *Hyg$^r$* | *prpR* deletion mutant with Hyg cassette inserted in the chromosomal copy of *prpR* gene (*Rv1129c*) | Kind gift from Christopher M. Sassetti |
| 4. | *ΔprpR::Hyg$^r$::lux* | *Hyg$^r$, Kan$^r$* | *prpR* marked deletion mutant with luciferase reporter | This work |
| 5. | *ΔmetZ* | | *Rv0391*, *metZ* deletion mutant, unmarked | This work |
| 6. | *ΔmetZ Comp* | *Kan$^r$* | *ΔmetZ* mutant harboring pST-K:*metZ* plasmid | This work |
| 7. | Rv::*lux* | *Zeo$^r$* | *Rv*-WT Luciferase reporter | This work |
| 8. | *ΔmetZ::lux* | *Zeo$^r$* | *ΔmetZ* Luciferase reporter | This work |
| 9. | *ΔmetZ Comp::lux* | *Kan$^r$, Zeo$^r$* | *ΔmetZ Comp* Luciferase reporter | This work |
| 10. | Rv: pMV261 mEmerald, TagRFP | *Hyg$^r$* | Mycobacterial Live Dead Reporter expressing Rv-WT | This work |
| 11. | *ΔmetZ*: pMV261 mEmerald, TagRFP | *Hyg$^r$* | Mycobacterial Live Dead Reporter expressing *ΔmetZ* mutant | This work |
| 12. | *ΔmetZ Comp*: pMV261 mEmerald, TagRFP | *Kan$^r$, Hyg$^r$* | Mycobacterial Live Dead Reporter expressing *ΔmetZ Comp* strain | This work |
| 13. | *Rv: Mrx1-roGFP2* | *Hyg$^r$* | Redox biosensor expressing Rv-WT | This work |
| 14. | *ΔmetZ: Mrx1-roGFP2* | *Hyg$^r$* | Redox biosensor expressing *ΔmetZ* mutant | This work |
| 15. | *ΔmetZ Comp*: *Mrx1-roGFP2* | *Hyg* | Redox biosensor expressing *ΔmetZ Comp* strain | This work |
| 16. | *Rv::metB$_{KD}$::lux* | *Kan$^r$* | Rv-WT lux reporter strain with *metB* gene knocked down using CRISPRi | This work |
| 17. | *ΔmetZ::metB$_{KD}$::lux* | *Kan$^r$* | *ΔmetZ* lux reporter strain with *metB* gene knocked down using CRISPRi | This work |
| 18. | *Rv::NT$_{KD}$* | *Kan$^r$* | Rv-WT with control sgRNA (non-targeting) CRISPRi control | This work |
