## Supplementary material for "pH-dependent direct sulfhydrylation pathway is required for the pathogenesis of *Mycobacterium tuberculosis*": List of Primers

**List of primers used in the study**

| S. No. | Primer Name | Sequence 5'-3' | Feature |
|---|---|---|---|
| 1. | *metZ*-F1 | GCTTAATTAAGTCGAGATCGAGTTTTTGGTCACC | Cloning of 1000bp upstream region of *metZ* gene for knockout construct |
| 2. | *metZ*-R1 | ATGCGGCCGCGCGGACCGAAGACTCGTC | |
| 3. | *metZ*-F2 | ATCTCGAGACCGACGACCTGATTGCCGAT | Cloning of 1000bp downstream region of *metZ* gene for knockout construct |
| 4. | *metZ*-R2 | ATCTCGAGCGGAATGAAGACCATCGACGAC | |
| 5. | *metZ*-F | AACATATGATGACCGACGAGTCTTCGGTC | Cloning of *metZ* encoding ORF in *NdeI/ EcoRI* sites |
| 6. | *metZ*-R | AAGAATTCTTAGCTCAACGCCCGATCGATATC | |
| 7. | *metB*-CR-F | GGGAACCGCGCCGGCGCCGTTCTG | Cloning of *metB* targeting sgRNA |
| 8. | *metB*-CR-R | AAACCAGAACGGCGCCGGCGCGGT | |
| 9. | *metB*-RT-F | GGCGGCACATTCCGGTTGATA | qPCR primers of *metB* |
| 10. | *metB*-RT-R | GTGAGGCAAAGGTATTGTCCACCA | |
| 11. | *metC*-RT-F | GACCATCTCCAACCCGCAG | qPCR primers of *metC* |
| 12. | *metC*-RT-R | GTCGGGGGTGGTGAAGC | |
| 13. | *metZ*-RT-F | GACGACCTCTCGCAATGGGA | qPCR primers of *metZ* |
| 14. | *metZ*-RT-R | GTACACCACCACGTCGACC | |
| 15. | *sigA*-RT-F | GAAGACCACGAAGACCTCGAAG | qPCR primers of *sigA* |
| 16. | *sigA*-RT-R | GTTTGAGGTAGGCGCGAACC | |
| 17. | *icl1*-RT-F | CAGCACATCCGCACTTTGAC | qPCR primers of *icl1* |
| 18. | *icl1*-RT-R | ATCACCACCGTGGGAACATC | |
